# The state-of-the-art machine learning model for Plasma Protein Binding Prediction: computational modeling with OCHEM and experimental validation

**DOI:** 10.1101/2024.07.12.603170

**Authors:** Zunsheng Han, Zhonghua Xia, Jie Xia, Igor V. Tetko, Song Wu

**Author notes:** corresponding authors Correspondence should be addressed to I.V.T. J.X. or S.W. Dr. Igor V. Tetko Institute of Structural Biology, Helmholtz Munich - German Research Center for Environmental Health, *Ingolstädter Landstraße 1, 85764 Neuherberg*, Germany *cBIGCHEM GmbH, Valerystr. 49, 85716 Unterschleißheim*, Germany Dr. Jie Xia and Song Wu Institute of Materia Medica, Chinese Academy of Medical Sciences, *No. 2 Nanwei Road*, Beijing 100050, China.

## Abstract

Plasma protein binding (PPB) is closely related to pharmacokinetics, pharmacodynamics and drug toxicity. Prediction of PPB is an alternative to experimental approaches that are known to be time-consuming and costly. Although there are various models and web servers for PPB prediction already available, they suffer from low prediction accuracy and poor interpretability, in particular for molecules with high values, and are most often not properly validated in prospective studies. Here, we carried out strict data curation, and applied consensus modeling to obtain a model with a coefficient of determination of 0.90 and 0.91 on the training set and the test set, respectively. This model was further validated in a prospective study to predict 63 poly-fluorinated and another 25 highly diverse compounds, and its performance for both these sets was superior to that of other previously reported models. To identify structural features related to PPB, we analyzed a model based on Morgan2 fingerprints and identified that features such as aromatic rings, halogen atoms, heterocyclic rings can discriminate high- and low-PPB molecules. In conclusion, we have established a PPB prediction model that showed state-of-the-art performance in prospective screening, which we have made publicly available in the OCHEM platform (https://ochem.eu/article/29).

**Graphic Abstract:** 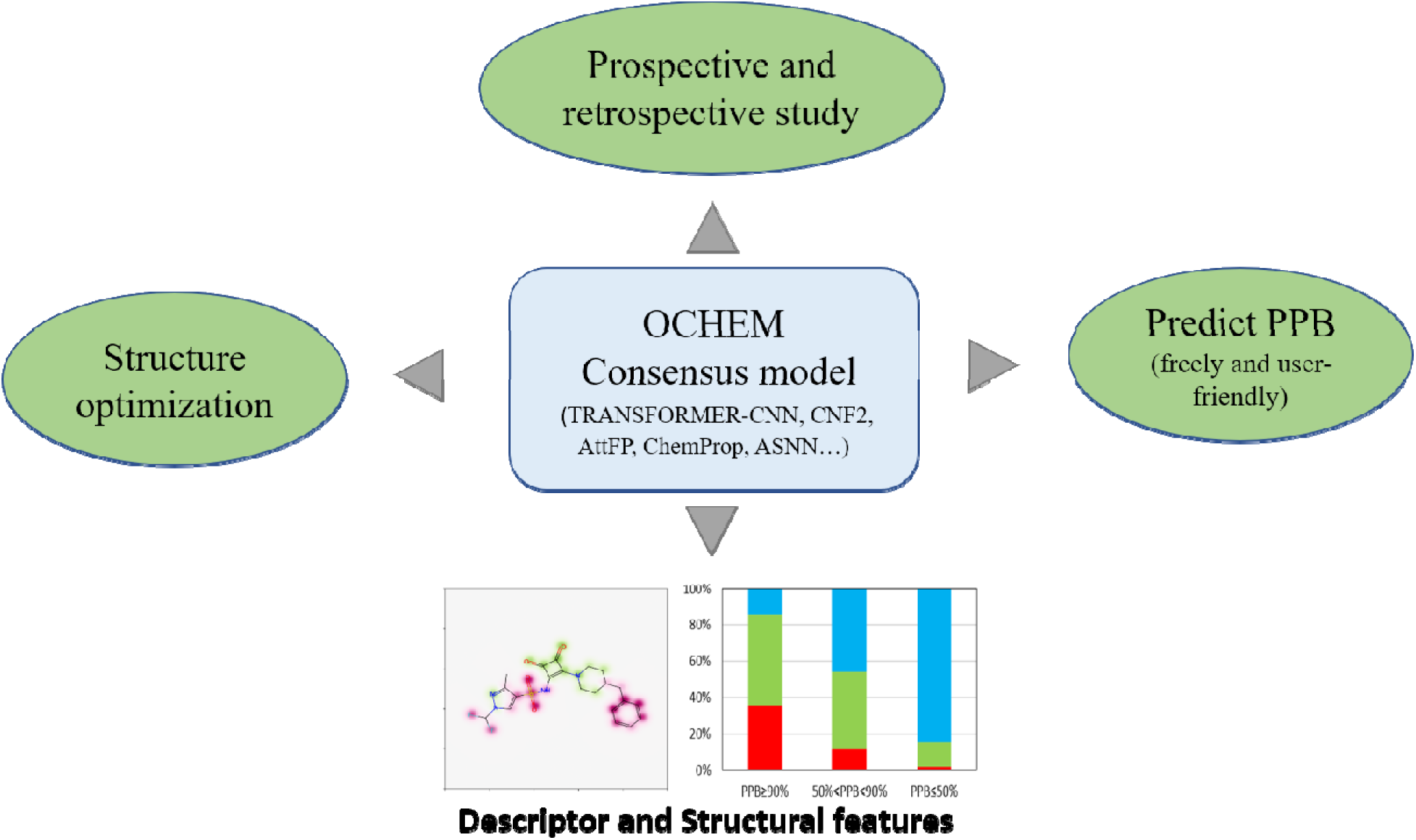

## 1. INTRODUCTION

Binding of drugs to plasma proteins is one of the important parameters of pharmacokinetics, as it can affect many key properties of drugs, *e.g.*, distribution volume (Vss), drug-drug interaction (DDI), clearance rate (CL) and therapeutic index (TI) ^[1-3]^. After drugs are administered, most drugs bind to plasma proteins to various degrees through blood circulation, forming drug-protein complexes. This binding is reversible and most of the time, it is non-specific ^[4, 5]^. Only free drugs (i.e., unbound drugs) can exert their biological effects. However, these drug-plasma protein complex can be used as a drug reservoir. When free drugs are metabolized and the concentration of free drugs is reduced, bound drugs can be released and become free drugs to play their role ^[4, 6]^. Drugs based on compounds with a high affinity to plasma proteins have an increased half-life, and in some cases higher doses of the drug may be required to achieve an effective concentration for treatment. In addition, drugs bind to plasma proteins competitively, and drugs with higher binding rates will occupy most of the plasma protein binding sites. Special attention should be paid to drug-drug interaction (DDI)^[1]^. Drugs with high PPB values may influence the binding of other drugs to the same plasma proteins, resulting in an increase or decrease in the (free) plasma concentration of the other drug, leading to toxicity or ineffectiveness. Situations like this mainly arise for drugs with narrow therapeutic windows, e.g., warfarin^[3, 7]^. Consequently, the PPB property of a drug affects its absorption, distribution, metabolism, elimination, and toxicity (ADME-T). Thus, assessment of PPB is very important for the development of new drugs and the safe use of clinical drugs The level of binding of a drug to plasma proteins is usually evaluated as PPB rate (PPB%) or free fraction (fu) ^[6, 8]^. There are three commonly used methods for determining PPB,, i.e., equilibrium dialysis (ED)^[9-11]^, ultrafiltration (UF)^[12, 13]^ and ultracentrifugation (UC)^[8, 14]^. The ED method is the gold standard and is often used as a reference for UF and UC. However, each of the three methods has its own advantages and disadvantages depending on the application ^[11]^. For non-specific compounds with good adsorption, UC is the first choice, but the cost is high. For compounds that are not stable in plasma, UF is preferred, but the non-specific adsorption is more serious. ED is generally applied to most compounds, but its drawbacks are obvious, such as changes in initial equilibrium state, non-specific binding, volume transformation, Donnan effect and protein leakage, all of which can influence results^[6, 10, 15]^. In any case, experimental assays are complex, time-consuming, and expensive, while *in silico* prediction has the advantages of being economical, simple, and fast, and able to facilitate rapid screening of a large number of compounds.

Over the past decade, significant advances have been made in ML based drug discovery ^[3, 16]^. In recent years, a number of regression models using ML have been built for PPB prediction. Table 1 summarizes some of the published models and performance metrics^[17-24]^. These models have made great progress in improving PPB prediction accuracy, but the performance of these models on test sets still needs some improvement. Moreover, all of the aforementioned models were not validated in prospective studies. It is also worth mentioning that none of the previous studies (with the exception of the study by Lou et al.^[24]^ using IDL-PPBopt) discussed the substructure and physicochemical properties that are highly related to PPB.

**Table 1.**
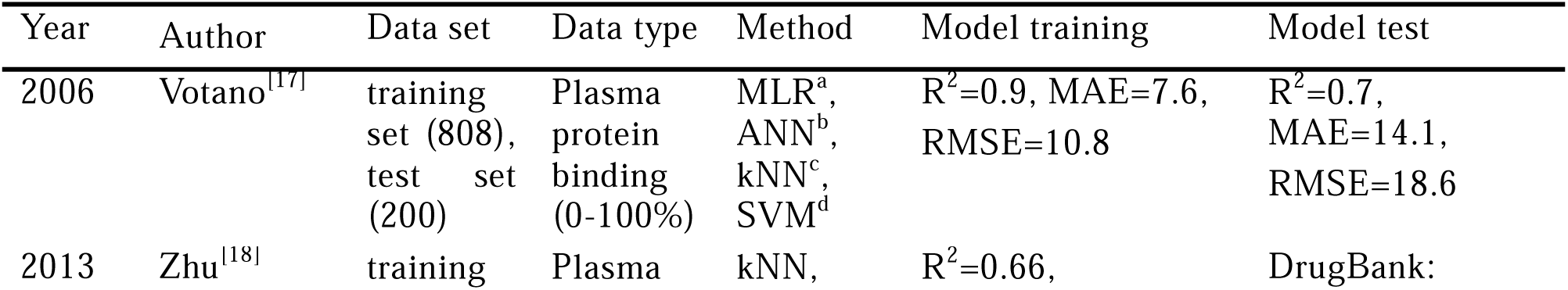
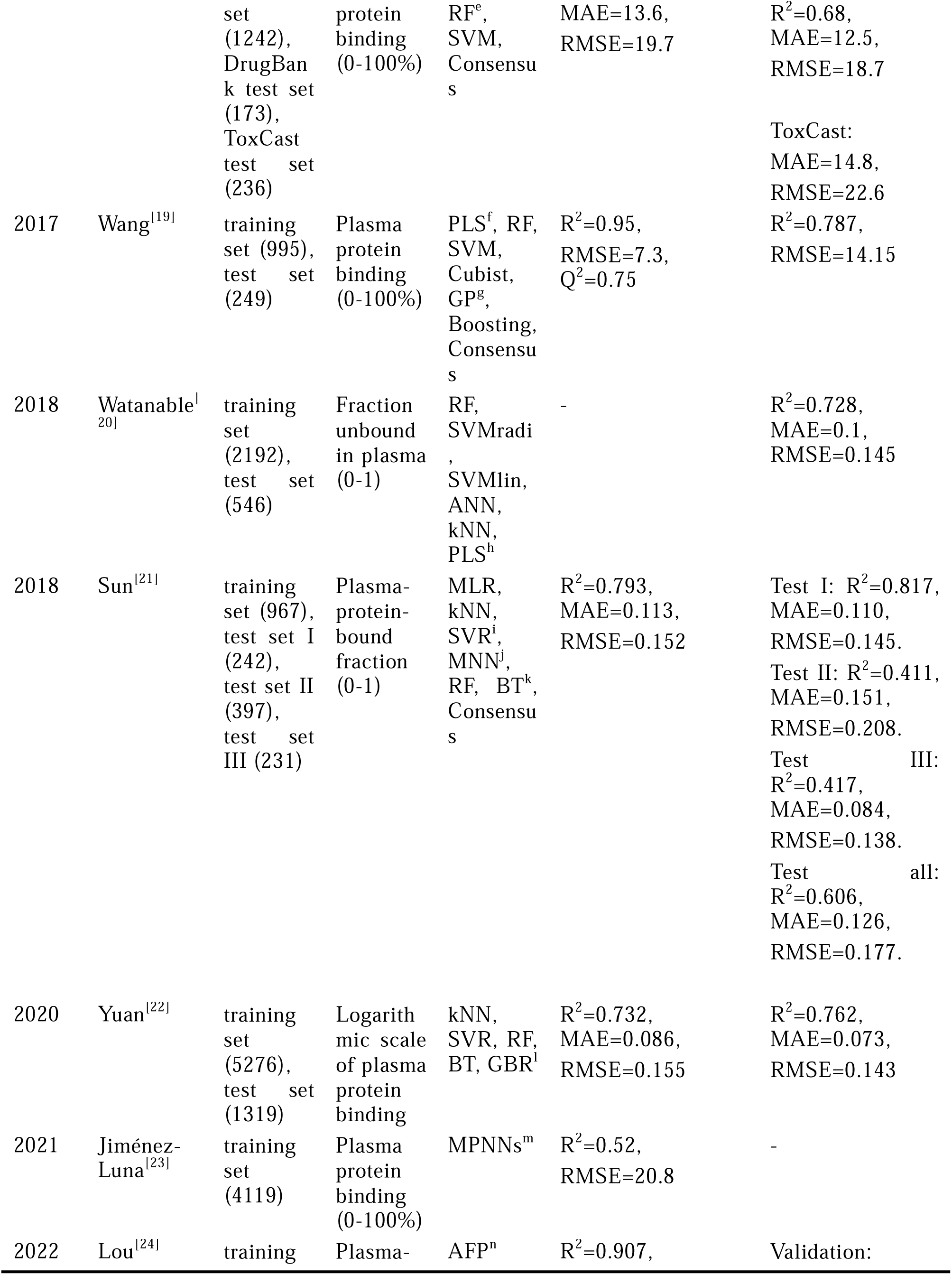
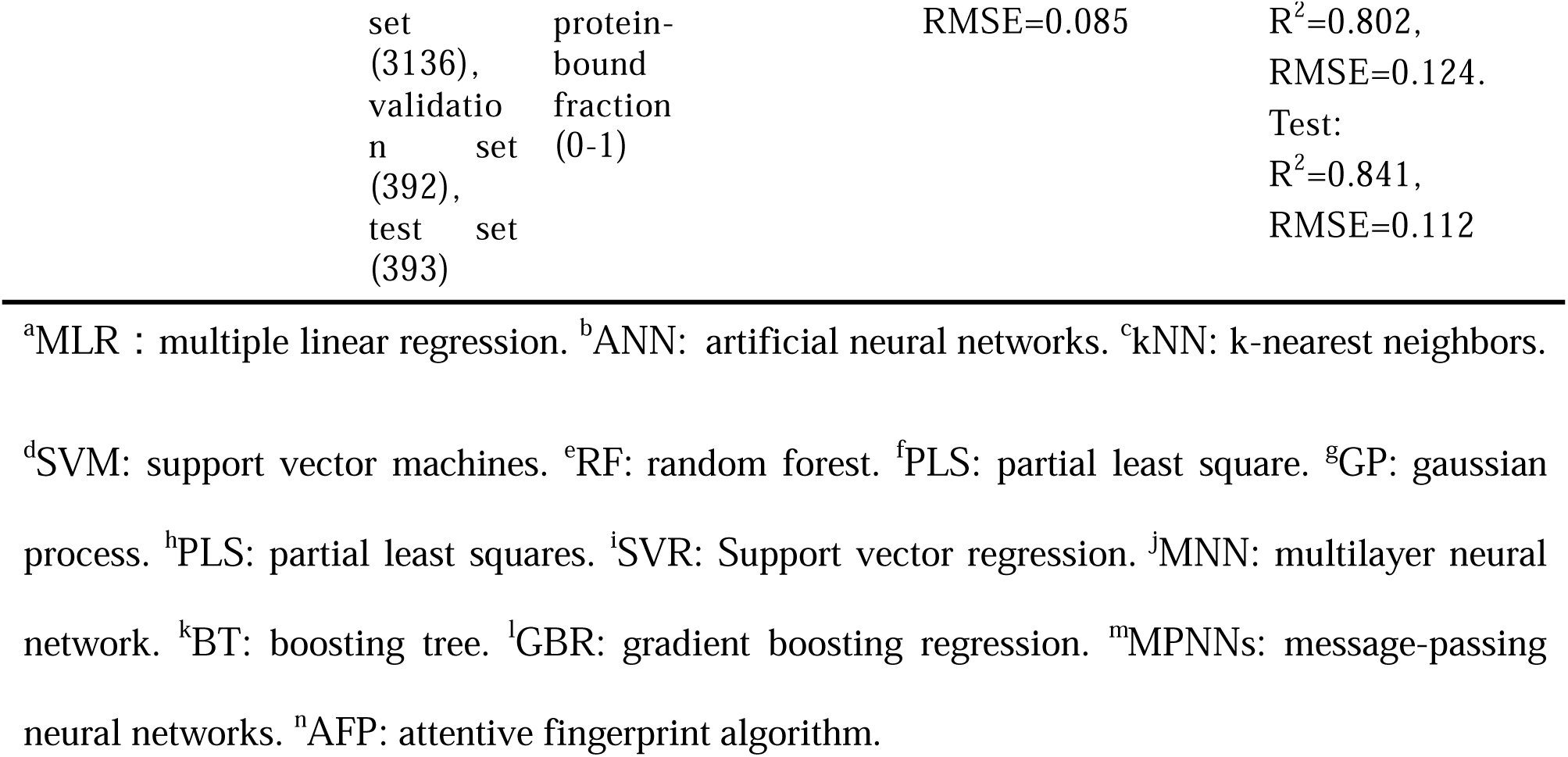
Performance of the Published Models.

In addition, the successful use of these models usually requires a significant amount of programming expertise, meaning that many of them are not particularly user-friendly. ^[25]^.To address this issue, several web servers offer free PPB prediction services, e.g. ADMETlab3.0^[26]^, admetSAR3.0^[27]^, DruMAP^[28]^, Pangu Drug^[29]^, pkCSM(Deep-PK)^[30]^, PreADMET^[31]^. Although these platforms are easy to use, it appears that the model accuracies of these platforms have not been validated in prospective studies.

On-line CHEmical database and Modelling environment (OCHEM) is an algorithm-rich, automated, and simple model training and sharing platform^[32]^. In this study, we leveraged the services offered by OCHEM to train models for PPB prediction with a variety of ML algorithms and descriptors. Then, we picked out several better-performing models and continued to train them into a consensus model. In addition, we performed experimental validation of the consensus model, by using it to predict the PPB of a diverse set of compounds from the ChemDiv library as an external validation set and then tested the compounds with ED assay. We also compared it with other publicly available web servers and models for such a prospective study of PPB prediction. Finally, we analyzed the molecular features that were associated with high and low PPB values. The workflow adopted in this study is shown in Figure 1.

**Figure 1.**
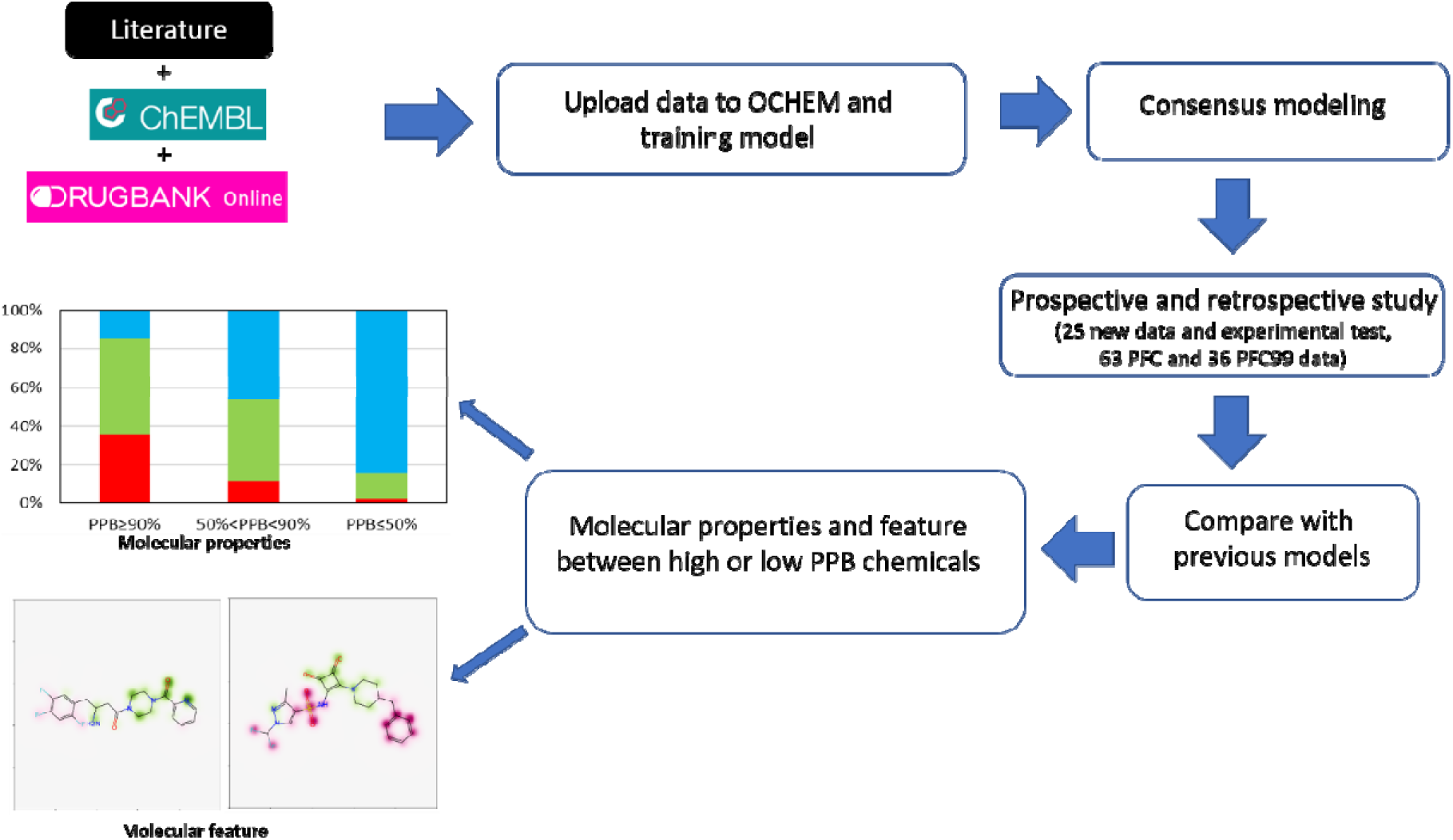
The workflow of the PPB prediction model.

## 2. Data

### 2.1 Training Data set

Human PPB data points were collected from several sources: (1) literature data from Tang^[24]^, Basant^[33]^, Reiko^[20]^, Brandall^[34]^, etc.; and (2) from DrugBank for drugs on market and in clinical trials (https://go.drugbank.com/, accessed in May 2022). A total of 9,500 data points were collected, including SMILES and multiple different forms of PPB values (e.g. protein binding rate 90% or 0.9 or unbound ratio 0.1 or 10%). The compound data were carefully prepared according to the following steps: (1) de-duplication, and removing mixtures, inorganic and organometallic compounds; (2) stripping salts and water; (3) removing compounds with molecular weights more than 800 Da and rotatable bonds more than 20 degrees^[35, 36]^.

According to the histogram of %PPB (cf. Figure S1), the distribution of original %PPB values was heavily skewed toward the high-PPB region. To minimize the effects of this skew, %PPB data points were transformed into a pseudo-equilibrium constant parameter (LogIt) in OCHEM automatically (see Eq. 1) as suggested elsewhere.^[37]^ After this conversion the distribution became Gaussian-like (cf. Figure 2A).

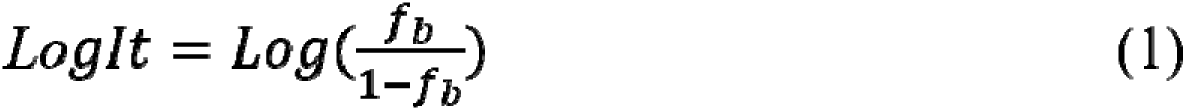

**Figure 2.**
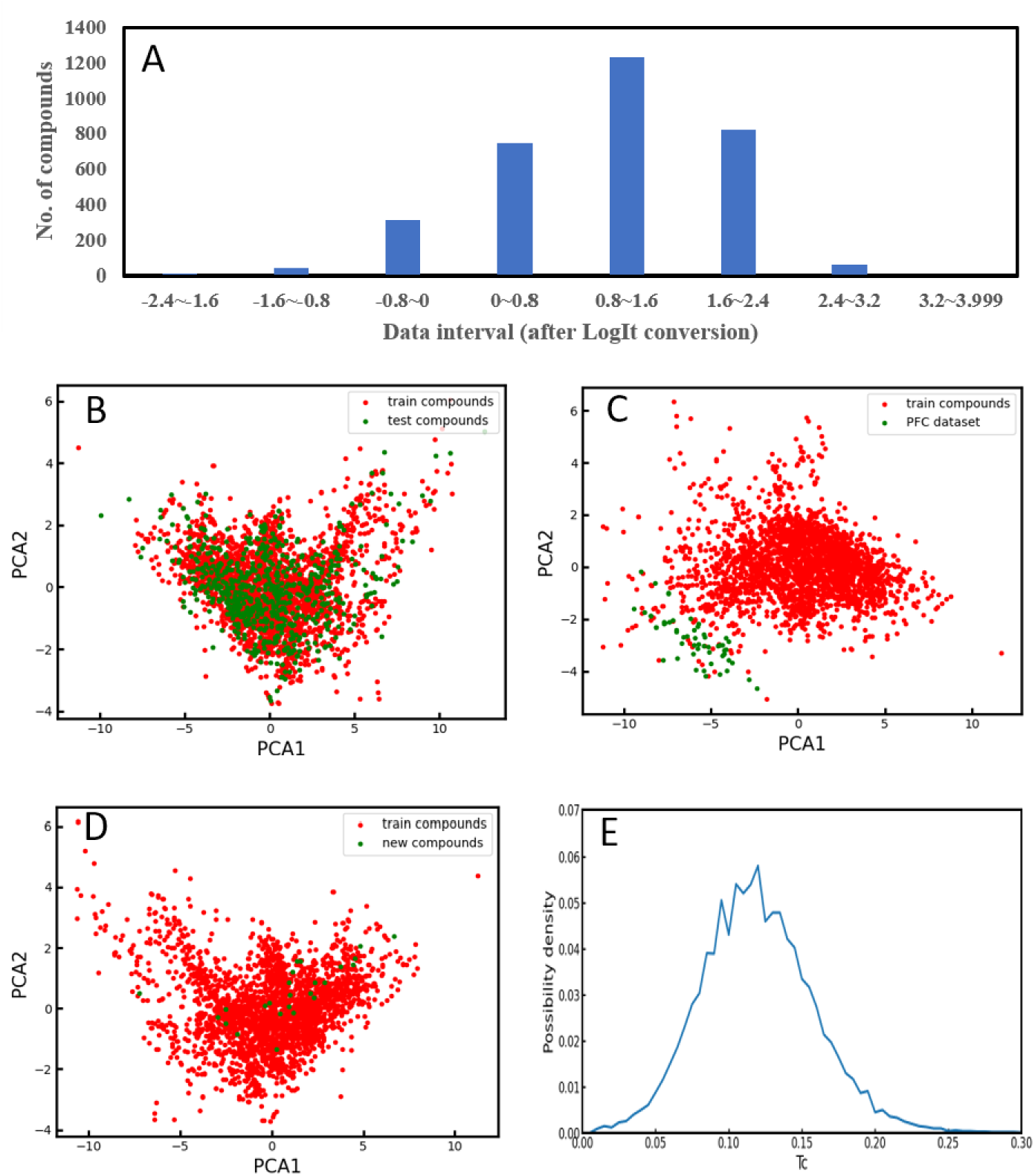
(A) Distribution of LogIt transformation dataset, PCA distribution plots: (B) Training and test dataset, (C) Training and PFC dataset, (D) Training and new dataset, (E) Pairwise Tanimoto coefficients distribution of 3,214 compounds.

where *f_b_* is fraction bound. This transformation is better for addressing/investigating differences in compounds with very small (i.e., < 1%) and very large (i.e., > 99%) PPB values. Indeed, there is a very large discrepancy in compounds with a PPB 99% and those with a PPB of 99.9%, since the difference in free concentration of the compound in plasma capable of producing a therapeutic effect will be 10 folds. Finally, total of 3214 compounds with PPB values were collected. These data were randomly split into training (2571) and external validation (643) sets.

### 2.2 Retrospective validation: PFC dataset

A dataset containing PPB values for Per- and Polyfluoroalkyl Substances was used for model validation^[38]^. As the article was published in 2023, this dataset was not yet encountered by many models published by other groups, nor by our own model, which collected data from earlier publications. We found that 5 out of 68 compounds from the novel dataset were also part of our training dataset. These molecules were excluded and the remaining 63 molecules were used as a validation set, which we called the “PFC dataset”. Furthermore, 36 compounds from the PFC set with PPB >99% formed the PFC99 set, which was used to compare methods for compounds that bind very strongly to plasma proteins.

### 2.3 Prospective validation: new dataset

A prospective study was performed to estimate model performance for a set of compounds selected using an experimental design method. The 10k compounds of the ChemDiv diverse subset purchased by Institute of Materia Medica, Chinese Academy of Medical Sciences (structure information can be retrieved at Zenodo: https://zenodo.org/records/12641856), were clustered using the “Cluster Ligands” module of Discovery Studio (version 2016). Specifically, the fixed “Number of Clusters” was set to 25 and FCFP_6 fingerprints^[39]^ were used to represent compounds. The cluster centers (molecules) were selected to form the external prospective set. These were all new compounds and there was no overlap between these molecules and the modeling set (training and retrospective test set).

The experimental method used to measure PPB was chosen based on the premise that free drugs can pass through the semi-permeable membrane (cellulose material), while the drugs bound to plasma proteins cannot. To an appropriate volume of compound stock solution, plasma (Human, IPHASE Pharmaceutical Technology Co., Ltd. Ethics No. ZYBFZ-LL-SOP-020-02) was added to yield a 10 μg/mL concentration of compound solution in plasma. 200 μL of compound solution in plasma was transferred to the dialysis bag of the RED device (Thermo Fisher Scientific, Single-Use RED Plate with Inserts, 8K MWCO), and a 400μL PBS solution was added to the corresponding well and the plate was covered. The RED dialysis unit was incubated at 37 °C in a constant-temperature incubator and shaken at 220 rpm for 4 h. After the incubation was completed, the acetonitrile protein precipitation method followed by HPLC or LC/MS/MS were used to determine the compound concentration in the dialysis bag and the compound concentration in the corresponding well. The results from the HPLC and LC/MS/MS experiments are shown in Table S1. The plasma protein binding rates PPB% were calculated according to the formula 100% - B/A * 100%.

For the chemical space analysis, the Mordred package(1.2.0)^[40]^ was used to calculate the descriptors, since it contributed the model with highest accuracy as shown in Results section. The Tanimoto coefficients (Tc)^[41]^ based on Morgan2 fingerprints (implementation of ECFP4^[39]^ in RDKit package^[42]^), were calculated for each pair of compounds in the initial set of 3,214 compounds and their distribution.

## 3 Methods

The PPB regression models were built by using the Online CHEmistry database and Modeling environment (OCHEM)^[32]^, a web server that integrates a wide variety of ML algorithms and molecular representations. During model training, a number of algorithms implemented in OCHEM were applied and the models giving the best performance were combined into a consensus model^[43]^ which provided the final predictions.

### 3.1 Analyzed machine approaches

We initially analyzed several machine learning approaches available in OCHEM, including traditional methods such as Partial Least Squares, Multiple Linear Regression Analysis, k-Nearest Neighbors as well as more advanced methods based on decision trees, such as Random Forest, XGboost, CatBoos, shallow and deep neural networks, which develop models based on calculated descriptors. Among all investigated methods, the Associative Neural Network (ASNN) method^[44]^ generally provided higher performances across analyzed descriptors and as such this method was selected for model development. The ASNN is a combination of an ensemble of single hidden layer neural networks and k Nearest Neighbors. It was inspired by thalamo-cortical organization^[45]^ of the brain and improves performance of ensemble by correcting its bias using errors of the most similar records from the training set. Of the representation learning methods we selected Transformer Convolutional Neural Networks (TransCNN^[46]^), along with its variation Transformer Convolutional Neural Fingerprint (CNF^[47]^) as well as two Graph Neural networks methods: AttentiveFP^[48]^ and ChemProp^[49]^, both of which are implemented as part of the KGCNN^[50]^ package in OCHEM.

### 3.2 Model validation

The models were validated in two steps. The first one was the five-fold cross-validation (5CV) based on the training set during model construction. During this step training dataset (n=2571) was randomly split into five non-overlapping parts. Four of five parts were used to develop the model and predict the remaining part, which was used as the validation set for the respective model. This procedure was repeated five times and predictions for individual validation sets were collected to form the prediction for the initial training set. The final model was developed with the whole initial training set according to the same procedure as was used for each individual model during the 5CV.

In addition to 5CV result the prediction performance of the final model was validated using prospective and retrospective test sets, which are described in the Data section.

### 3.3 Descriptor filtering

OCHEM contains around 20 types of descriptors. For this analysis we decided to use only 2D descriptors. Indeed, we noticed that 3D based descriptors did not increase quality of models, e.g. models developed with full and 2D sets of Mordred^[40]^ descriptors had a similar accuracy, which is why we decided to proceed with 2D descriptors.

On each validation fold, as well as during the final model development, an unsupervised filtering of descriptors was performed. The pairwise de-correlation cutoff of descriptors was set to 0.95, and descriptors with low variance and nearly constant values were eliminated. The filtering procedure and descriptor and model hyperparameters were used with default values as specified in OCHEM.

### 3.4 Assessment of Model Performance

Three statistical parameters were used to evaluate model performance (cf. Eqs. 2-4). MAE and RMSE were used to evaluate the difference between the predicted values and the observed values. In addition to these parameters, the coefficient of determination R^2^, which measures explained variance of the model, was also used. A good model usually has small MAE and RMSE values, and an R^2^ value close to 1.

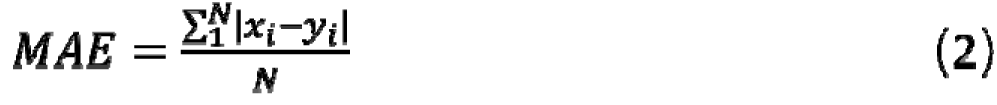

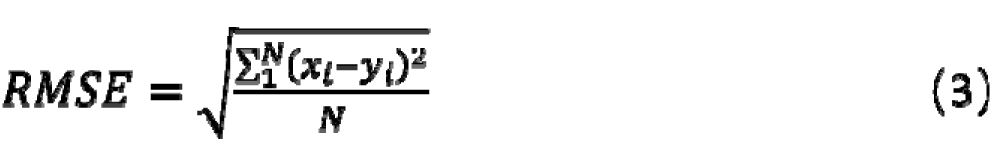

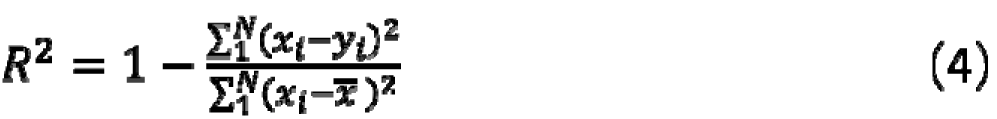

where is the observed value, is the predicted value, ̅ is the average of the observed values, *N* is the number of data points.

### 3.5 Applicability domain

The application domain (AD) is an important concept in quantitative structure activity relationships (QSAR)^[51]^. The measurement of AD was based on the distance to model (DM) of the compound, where DM is a numerical measure proportional to the model’s prediction uncertainty for a given compound ^[52]^. A large DM value corresponds to a low prediction accuracy for the target compound. In our study, the DM corresponds to the standard deviation of models in consensus. A DM that covered 95% of compounds in the training set was used to define the AD of a model. Each prediction was given a confidence interval to help users judge the reliability of the prediction.

### 3.6 PPB prediction using webservers

The PPB values for retrospective and prospective test set molecules were predicted using different computational platforms (ADMETlab3.0^[26]^ (https://admetlab3.scbdd.com), admetSAR3.0^[27]^ (http://lmmd.ecust.edu.cn/admetsar3), DruMAP^[28]^(https://drumap.nibiohn.go.jp), Pangu Drug^[29]^ (http://pangu-drug.com), PreADMET^[31]^ (https://preadmet.qsarhub.com/), pkCSM(Deep-PK)^[30]^(https://biosig.lab.uq.edu.au/deeppk/data). The prediction accuracies of these platforms were compared with performance of the PPB model developed in this study.

### 3.7 Feature analysis related to PPB

#### 3.7.1 Physicochemical descriptors analysis

At first, the physicochemical descriptors of all compounds (3,214) were calculated using the Mordred package (1.2.0)^[40]^, and then the Person correlation coefficient (*r*^2^) of each descriptor with PPB was calculated. Lastly, top 17 descriptors with *r^2^>0.4* were used to plot the heat map.

In order to gain insight into the model’s interpretability, the relationship between each of the aforementioned descriptors with high, medium and low PPB values were analyzed, respectively. All compounds were divided into three categories according to their PPB values, i.e. low PPB class (≤ 50%), medium PPB class (50-90%), high PPB class (≥90%). In each class, the compounds were further divided into 3 groups according to the descriptor being studied. Then, the number of compounds in each descriptor group in the categories of low, medium and high PPB was counted. The number of compounds in a certain category was defined as 100%, and the percentage of molecules in each descriptor group in all compounds in this category was calculated. The data were converted into graphs for visualization after calculation. Taking the descriptor of SlogP as an example, the high PPB compounds were divided into three subgroups (SLogP≥3, 1<SLogP<3, SLogP≤1), and the proportion of compounds in each subgroup was determined. This facilitated the identification of physicochemical properties associated with high and low-PPB compounds.

#### 3.7.2 Privileged substructures analysis with Similarity Map and Setcompare in OCHEM

A PPB classification model was built and was used to produce a similarity map.^[53]^ Specifically, the compounds from the initial set (n=3128) were divided into high and low PPB sets with thresholds of PPB >90% and PPB<50% respectively. The compounds with PPB values between 50% and 90% were removed. Each compound was represented with Morgan2 fingerprints. The classification model was built based on the training set and was evaluated on the test set. In the similarity map^[53]^, the atoms of each compound were marked with different colors according to the contribution value of the atom. The substructures composed of atoms with positive contributions or negative contributions were visualized.

The high/low PPB data were also analyzed using the SetCompare utility^[54]^ in combination with the Extended Functional Groups^[55]^ descriptor type. SetCompare uses hypergeometric distribution with Bonferroni correction to identify overrepresented descriptors amid compounds with high/low PPB data.

## 4. Results and discussion

### 4.1 The Data Set

After careful pre-processing, 3,214 PPB data points were obtained. The LogIt transformed data exhibited a Gaussian-like distribution, as shown in Figure 2(A). As mentioned in the Data section, the data points were randomly split between the training set and test set. The chemical space of the training, test, PFC and new compound datasets are shown in Figure 2(B-D) based on principal component analysis (PCA) of the 17 Mordred descriptors (see Supporting Material). The scatter plot shows substantial overlap in chemical space between training and test compounds. The Tanimoto coefficient (Tc) values of most pairs of compounds were less than 0.2 (see Figure 2(E)), indicating high chemical diversity in the PPB data set.

### 4.2 Models constructed in OCHEM

OCHEM offers a number of ML algorithms and various molecular representations. Almost all of the ML algorithms implemented in the platform were tested in this study. ASNN^[44]^ provided on average better performance compared to other descriptor-bases machine learning methods and were used for further analysis. Amid ASNN models, the model based on Mordred descriptors achieved the highest accuracy for the training set (Table 2). After comparing all models, we selected those with RMSE equal or less than 0.33 to build the consensus model. There were five ASNN models developed with ALogPS-Oestate^[56-58]^, EPA (https://www.epa.gov/comptox-tools/toxicity-estimation-software-tool-test), Fragmentor^[59]^, MOLD2^[60]^ and MORDRED (2D)^[40]^ descriptors as well as four models based on representation learning: transformer convolutional neural networks (Transformer CNN)^[61]^, Convolutional neural network Fingerprint (CNF2)^[62]^, ChemProp^[63]^ and AttFP^[64]^ (as implemented in KGCNN^[50]^ package). The performance of the consensus model, calculated as a simple average of these selected models, was superior to any individual model for both training and external test sets (see Table 2 and Figure 3).

**Figure 3.**
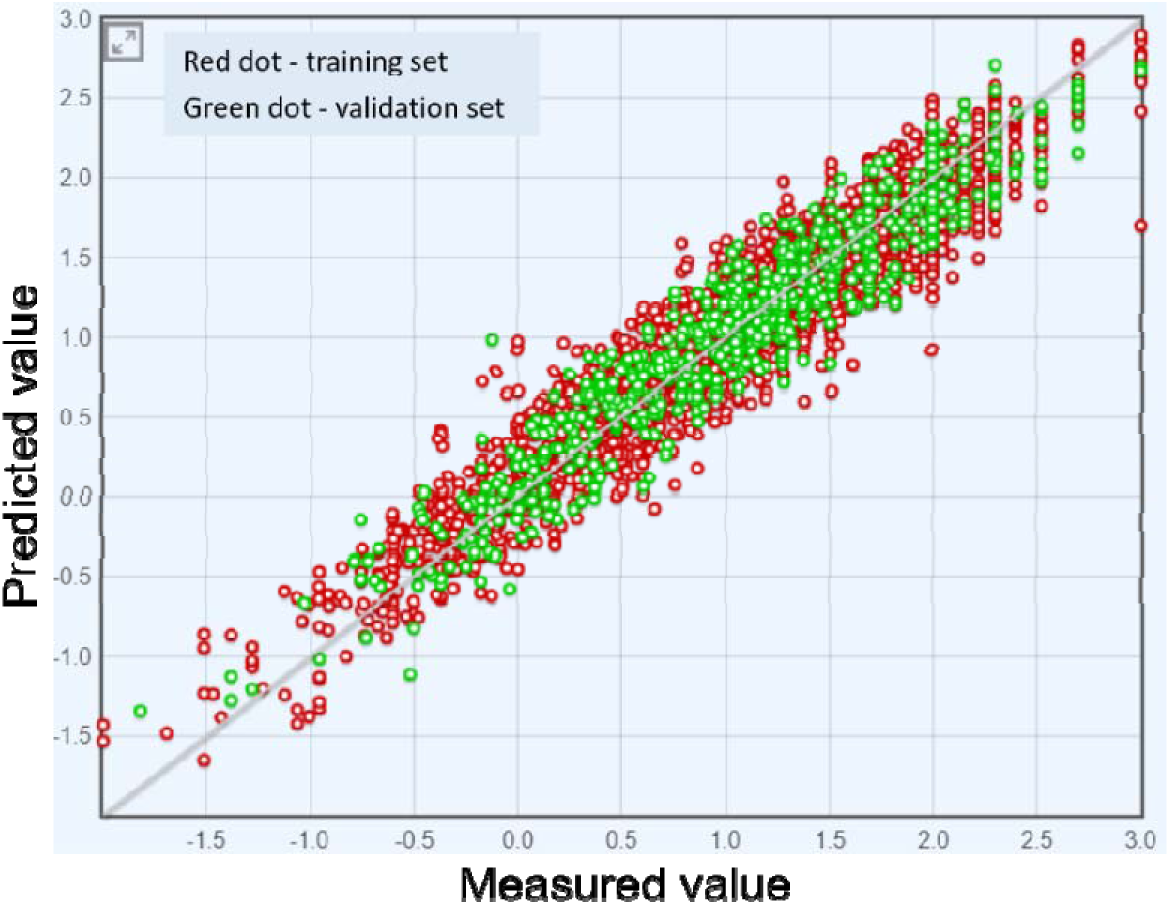
The measured PPB vs the PPB predicted by the consensus model for the training and test sets.

**Table 2.**
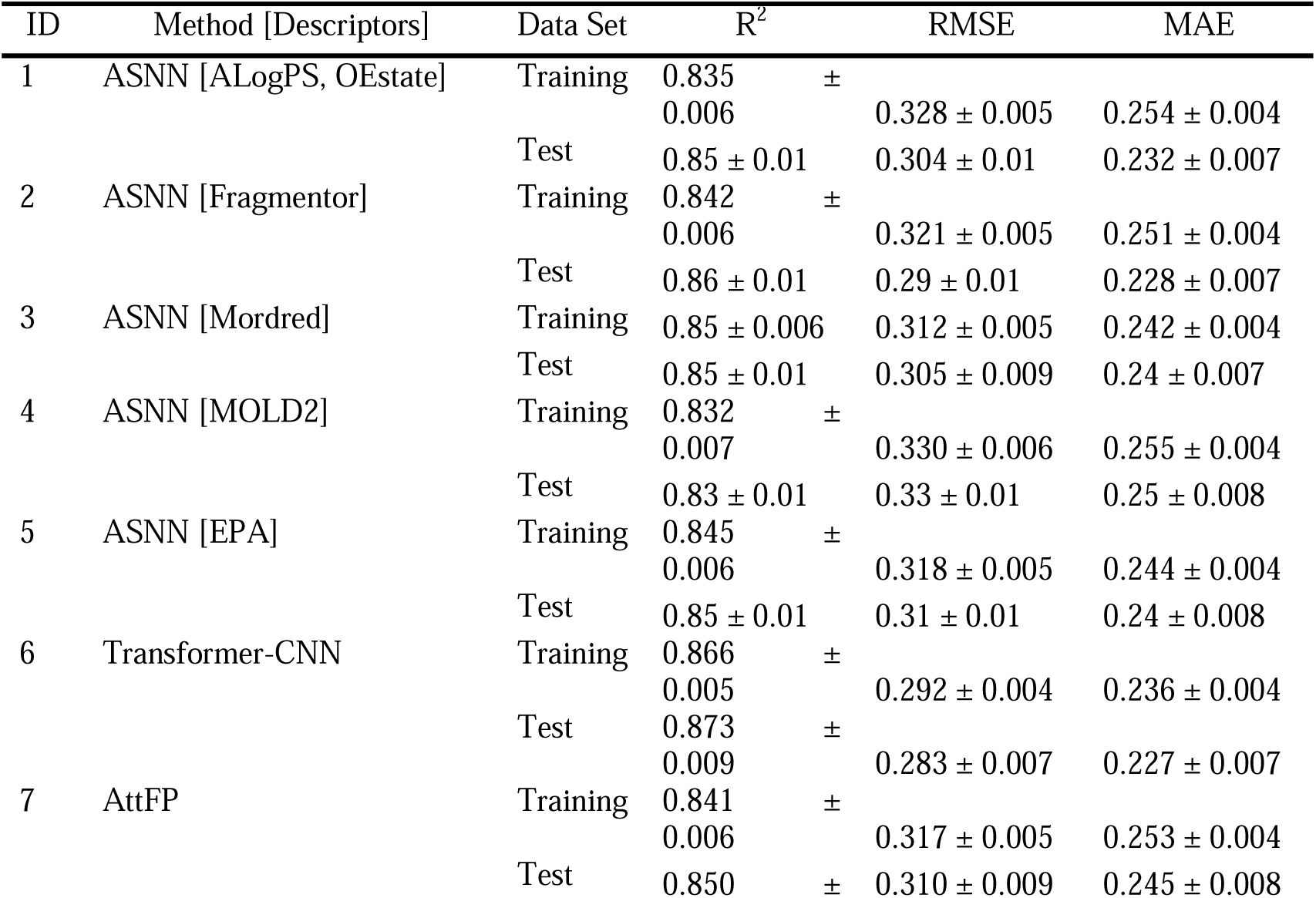
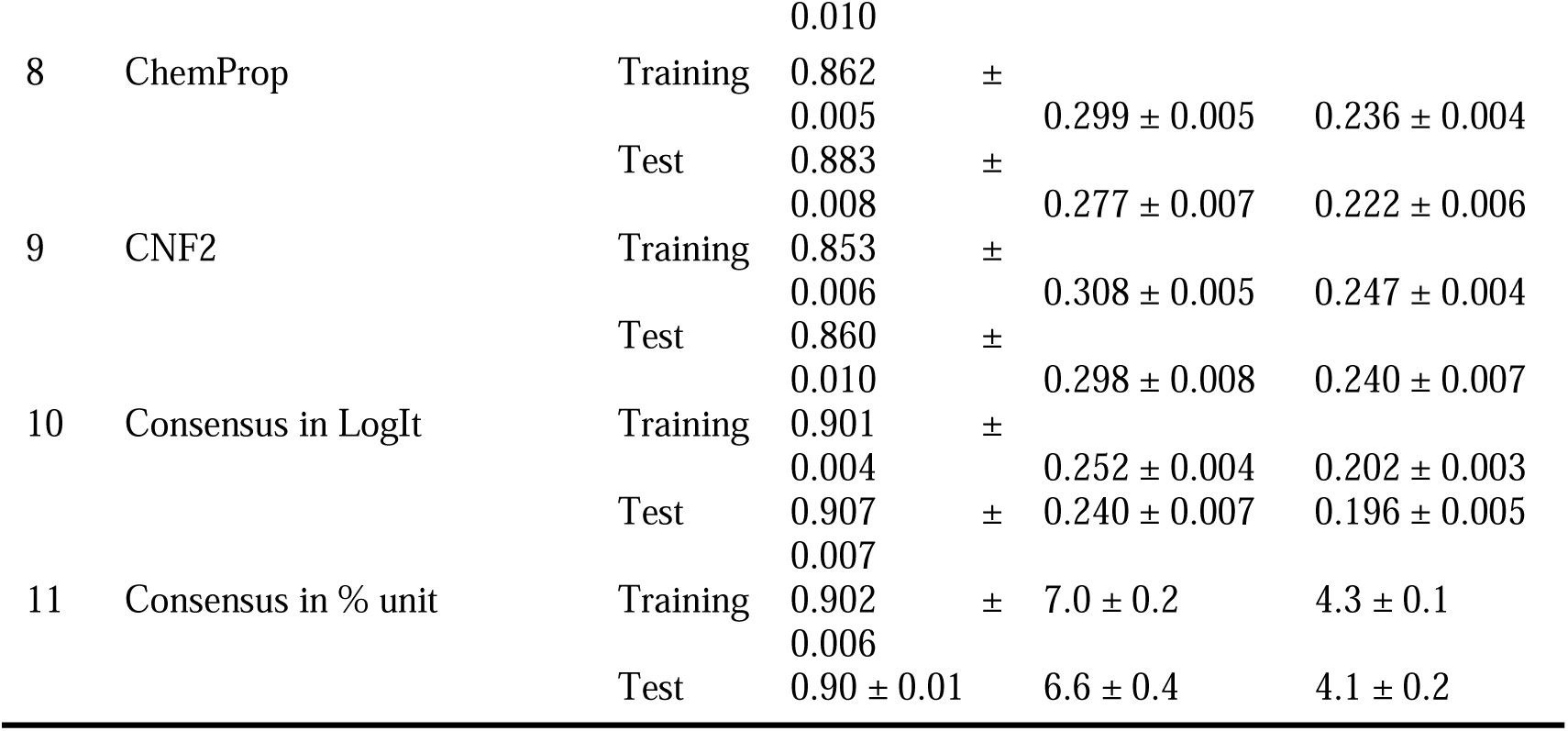
Performance of individual and the consensus models.

### 4.3 Prospective and retrospective study: Model performance and comparison with other models

We clustered a diverse ChemDiv set (10,000) based on FCFP_6 fingerprints into 25 clusters and selected 25 cluster centers to comprise an external set for the prospective study. The PPB of the compounds were predicted using several representative open platforms, including ADMETlab3.0, admetSAR3.0, DruMAP, Pangu Drug, pkCSM(Deep-PK), PreADMET and the newly developed consensus model in OCHEM. To compare the prediction accuracy of these models in practice, we determined the true PPB values of 25 new compounds using equilibrium dialysis combined with HPLC or LC-MS/MS. The structures and measured PPB values of the 25 compounds are shown in Figure 4. The measured values and results of each prediction platform are shown in Supporting Material. The predictive performance parameters for the different platforms are listed in Table 3. The consensus model developed in this study achieved a higher accuracy than the other published models.

**Figure 4.**
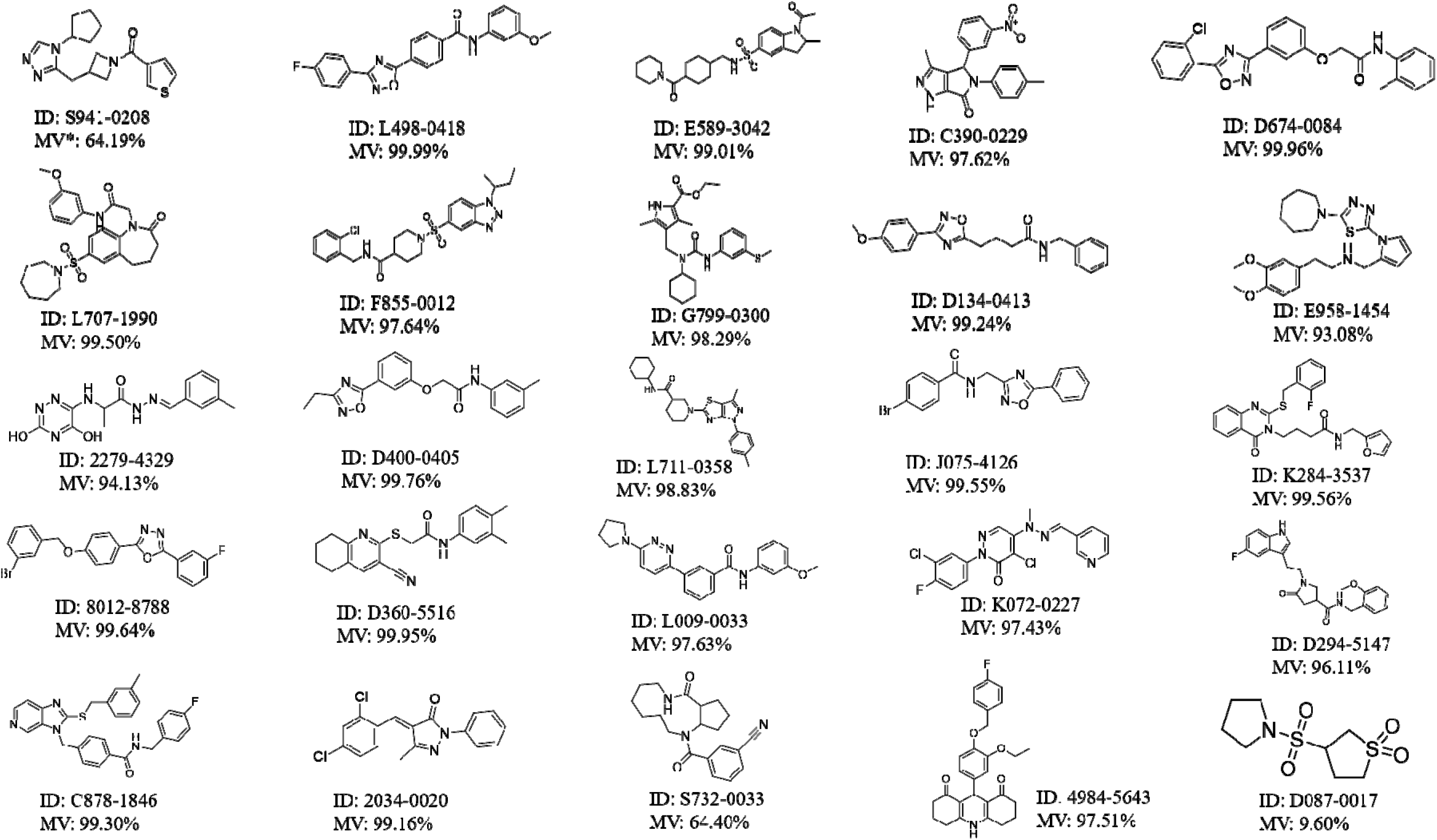
The structure and experimental results of the 25 new compounds. (*MV: measured value).

**Table 3.**
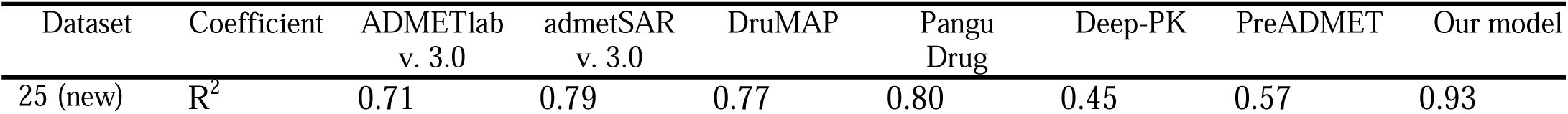
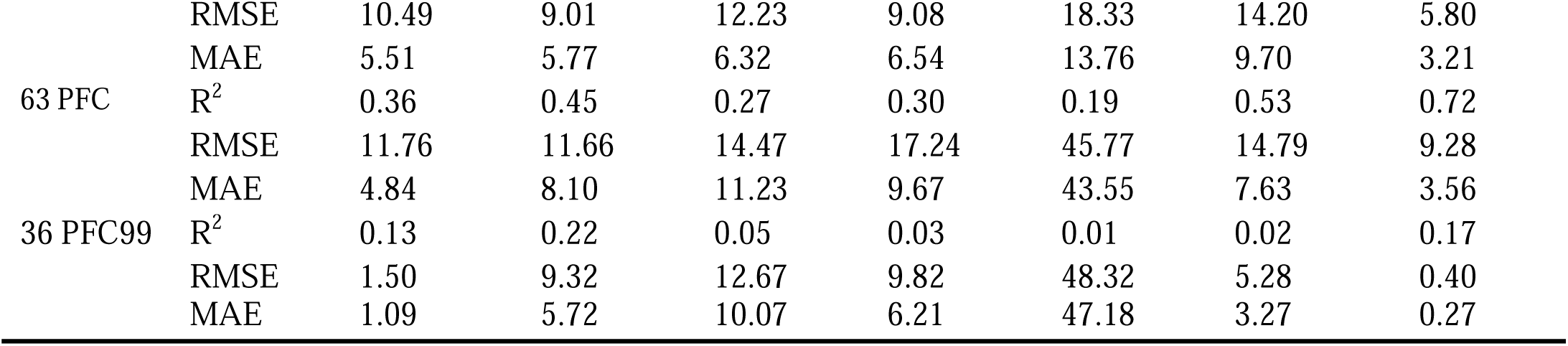
The tested platforms’ prediction performance for external validation sets.

For the retrospective study, we also compared the predictive performances of the models with the 63 PFC dataset (See Table 3 and Supporting Material for more detail) and found that the consensus model developed in this study also achieved a higher accuracy for this set than the other platforms. In addition, the results for compounds with PPB>99% (PFC99 set) predicted by the developed model had significantly better RMSE values (0.4%) compared to the results obtained by other models, which had RMSE values 3 to 100 times higher for the same compounds. Accurately predicting compounds with PPB>99% is very important and significant for drug discovery projects, because for binding rates of 99%, 99.9% or 99.99%, the free drug concentrations will differ by a factor of 10 or 100. Thus, despite the fact that existing models have a relatively good overall prediction performance when using %PPB as an overall performance measure, most of them were not able to distinguish compounds with very high PPBs.

### 4.4 Importance of LogIt function for predicting compounds with high PPB values

We used LogIt units to develop the individual and consensus models and estimate their performances. Of course, one can use different units, e.g. percentage, to estimate performance of the developed models by converting predicted and experimental values to the respective unit. For the consensus model we first converted LogIt to a percentage (%) for each individual model and then built a consensus by averaging predictions of individual models given in percentages. This was done in OCHEM, which allow the user to select a different target unit when creating a consensus model. As shown in Table 2 and Table 4, this conversion had a generally negative impact on the models: it decreased *R^2^* and also led to widened confidence intervals for RMSE and MAE. Therefore, we also developed individual models using the % unit and created a consensus model with % or LogIt units, respectively (See Table 4).

**Table 4.** RMSE of consensus models developed with LogIt and percentage units.

| Unit of individual models | Consensus model unit | Model name | Training set CV, n=2571 | Test set, n=643 | New set n=25 | Test set PFC, n=63 | Test set PFC99, n=36 |
| --- | --- | --- | --- | --- | --- | --- | --- |
| LogIt | LogIt | LogIt-LogIt | 0.252±0.004* | 0.24±0.007* | 0.68±0.1* | 0.79±0.08 | 0.49±0.05* |
|  | % | LogIt-% | 7±0.2% | 6.6±0.4% | 6±2% | 9±3 | 0.5±0.1%* |
| % | LogIt | %-LogIt | 0.293±0.004 | 0.273±0.008 | 0.78±0.1 | 0.74±0.07 | 0.9±0.1 |
|  | % | %-% | 7.1±0.2% | 6.4±0.4% | 7.1±2% | 7.5±1 | 3.6±0.9% |

PFC99 – dataset comprising compounds with PPB>99%. \*indicates significantly lower RMSE ( $p<0.05$ ) using the respective consensus unit.

The consensus model based on individual models developed with the same unit (LogIt-Logit model) had a significantly lower RMSE compared to the consensus model based on the individual models developed with the % unit (%-LogIt model) for all but the PFC set. For the latter set, RMSEs of both consensus models were not significantly different due to their large confidence intervals. There were no significant differences for all but the PFC99 subset (see discussion below) when we compared performances of LogIt-% and %-% models using the % unit.

The results for the PFC set had significantly large errors (RMSE=0.79 and 0.74 in LogIt unit) compared to the training set compounds (RMSE=0.25 and 0.29) for LogIt-LogIt and %-LogIt models, thus indicating that this set was particularly difficult to predict. This was the expected results, since the distribution of the compounds in the PFC set was markedly different to that of the training set, as shown on PCA plot (Figure 2C). However, there were no significant differences in RMSEs for the training and test sets when predicted values were compared using % units. However, for the subset of the PFC set with PPB>99% (PFC99, n = 36), the consensus model LogIt-* models yielded significantly smaller errors than the %-* consensus model for both unit types (Table 4). Thus, the LogIt-* consensus models were much better at predicting the PPB of compounds with very high binding. The ability to differentiate compounds with high PPB values is important for drug discovery, and we have shown that developing models using LogIt units allows for a deeper exploration of the data and more accurate predictions of PPB.

### 4.5 Physicochemical descriptors highly related to PPB

We calculated the physicochemical descriptors of the molecules, along with their corresponding Person correlation coefficients *r*^2^ with PPB using the Mordred program^[40]^. A total of 397 related physicochemical descriptors (*r*^2^ > 0.05) were obtained. Among them, 17 descriptors (see Supporting Material Table S2) were highly correlated to PPB, with *r^2^* values greater than 0.4. The heatmap below (Figure 5) illustrates the correlations between the selected descriptors and PPB.

**Figure 5.**
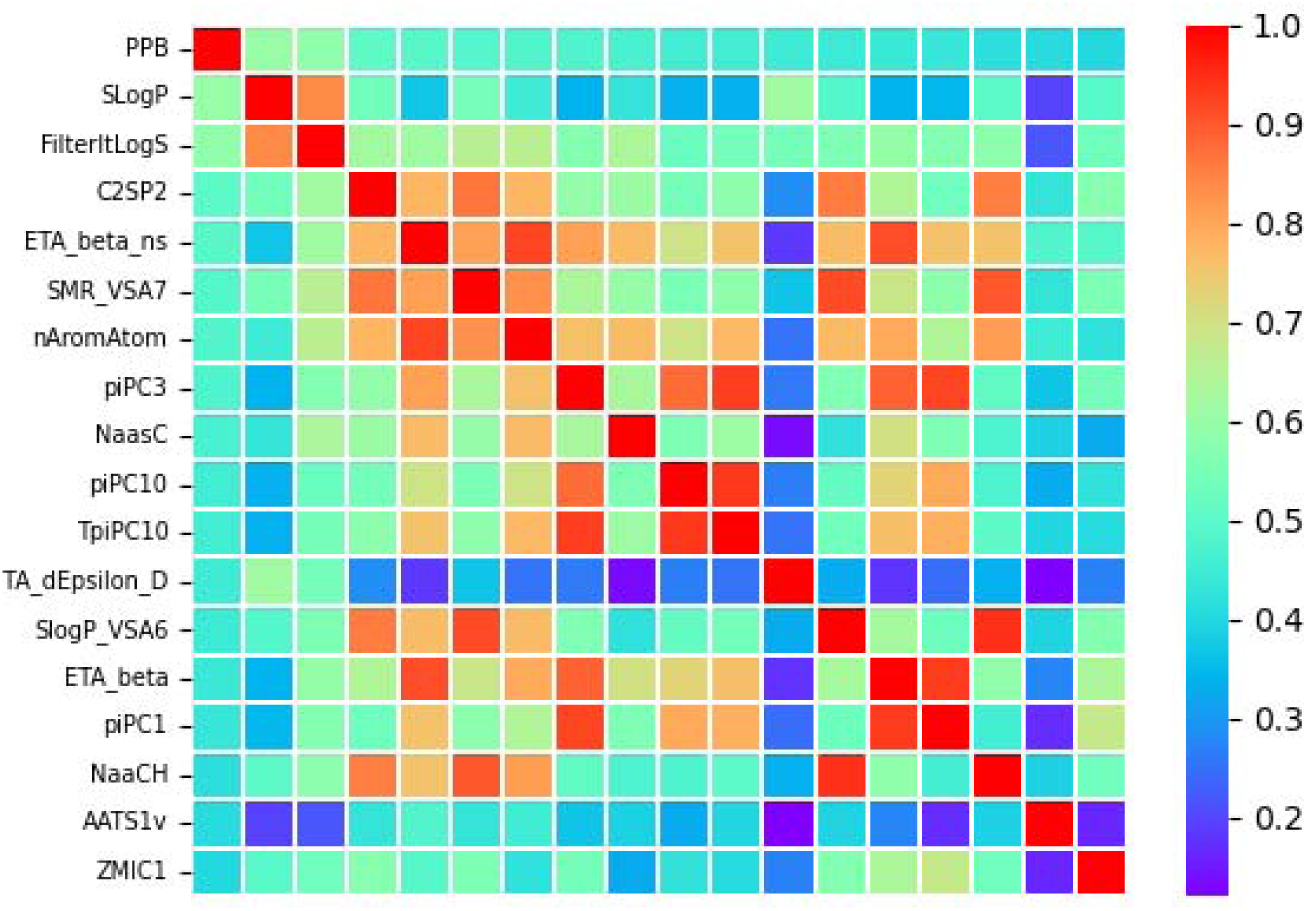
A heatmap of *r*^2^ between any two descriptors or between any descriptor and PPB. The heatmap was plotted based on the absolute value of *r*^2^.

As mentioned in the Methods section, we analyzed the relationships between each of the aforementioned descriptors and high, medium and low PPB values. We observed that the SLogP and LogS are very important physicochemical indicators for PPB, with R^2^ values are greater than 0.6. When considering three subgroups of SLogP (SLogP≥3, 1<SLog P<3, SLog P≤1) and three subgroups of LogS (LogS<-6, -6≤logS<-4, -4≤logS), we observe that a large proportion (around 80%) of high-PPB compounds had high SLogP (≥3) and low LogS (<-4), as shown in Figure 6A and 6B. In medium- or low-PPB compounds, the proportion of compounds with SLogP≥3 and LogS<-4 decreased significantly. Therefore, SLogP was positively correlated with PPB and LogS were negatively correlated with PPB.

**Figure 6.**
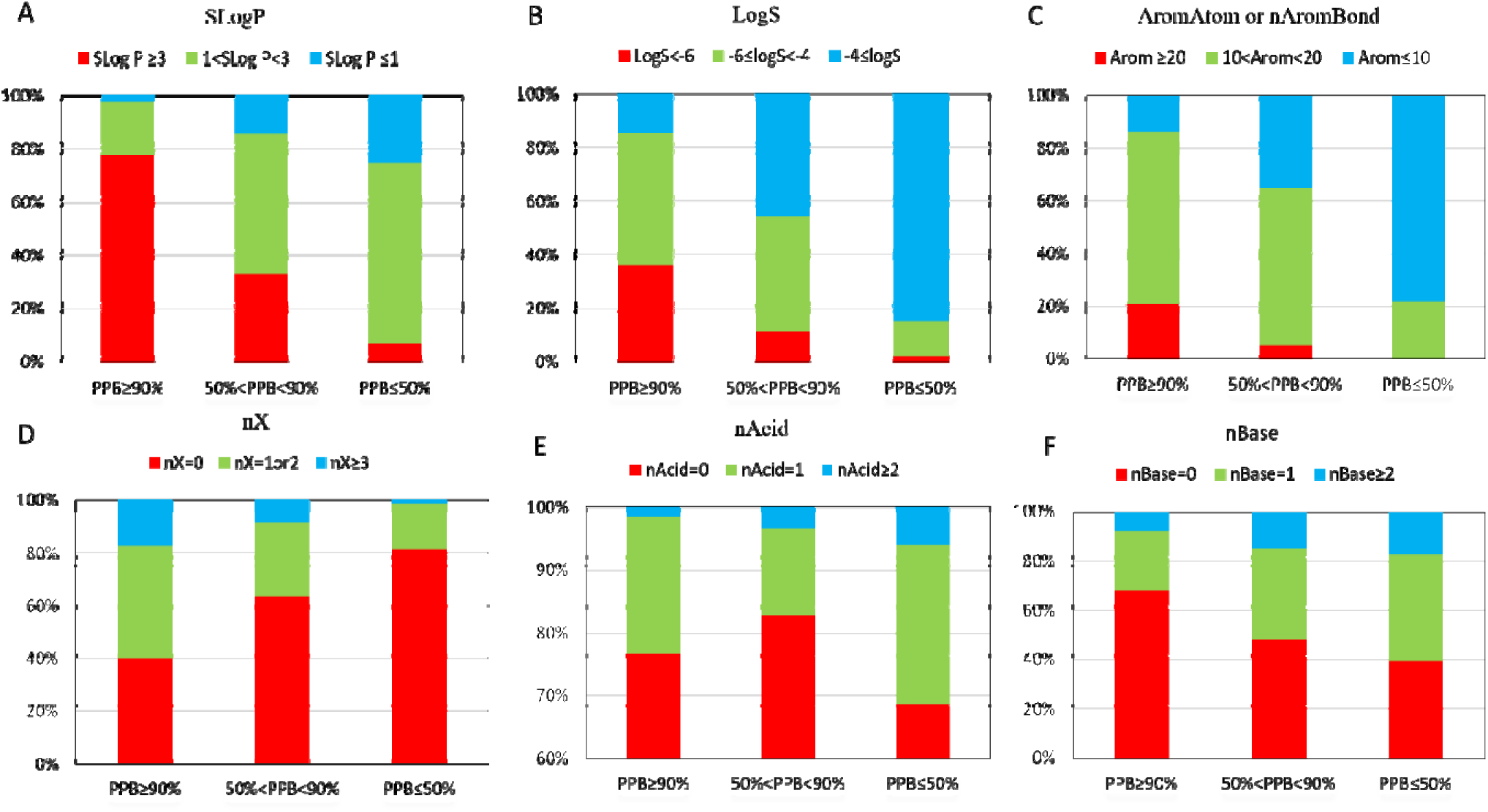
The differences in physicochemical properties of high-, medium- and low-PPB compounds. (A) SLogP, (B) LogS, (C) AromAtom or nAromBond, (D) nX, (E) nAcid, (F) nBas.

The number of AromAtom or nAromBond was also correlated with PPB (see Figure 6C.). A reduction in the number of AromAtom or nAromBond led to a significant decrease in the value of PPB – not a surprising observation, as it is generally known that the increase in the number of aromatic rings can improve the fat-soluble properties of a drug. In addition, we found that a higher number halogen atom (nX) may be associated with higher PPB values (Figure 6D).

The number of acid (nAcid) and basic (nBase) groups is closely related to the pKa of compounds. It was found that, for compounds with 0-1 acid group, nAcid had no influence on PPB. For nAcid ≤ 2, the increase of nAcid may lead to lower PPB values. As for nBase, the effect of basic groups on PPB was more significant than that of acidic groups. In general, the higher the number of basic groups, the lower the PPB value (Figure 6E and 6F).

Atom-bond Connectivity Index (ABC Index) refers to the strong interaction between two or more adjacent atoms, including covalent bond,s ionic bonds and metallic bonds. The study found that lower ABC Indices were associated with lower PPB values, as shown in Supporting Material Figure S2A. Hydrogen bonding is known to be a very important intermolecular force, and our results showed that the more hydrogen bond donors a molecule contained (i.e. >2), the more likely it was to have a lower PPB, whereas the number of hydrogen bond receptors had no significant influence on PPB (see Supporting Material Figure S2B and S2C). Furthermore, we found that some feature descriptors also showed some positive correlation with PPB. For example: KappaShapeIndex, sum of atomic volume parameters (McGowanVolume), atomic polarizability, rotatable bond (nRot), van der Waals volume (VdwVolumeABC), and some topological indicators, such as Wiener index and ZagrebIndex, see Supporting Material Figure S2D-J.

### 4.6 Substructures that affect PPB

To identify structures that significantly influence PPB, we built a classification model (PPB>90% and PPB< 50%) using Morgan2 fingerprints and generated a similarity map for several representative compounds^[53]^. The representative similarity maps of 3 low-PPB and 3 high-PPB compounds are displayed in Figure 7. In low-PPB compounds (see Figure 7A-C), we can conclude that: (1) amino groups often appeared in low-PPB molecules, and secondary and primary amines were dominant, which was exactly consistent with the conclusion of the descriptor analysis (the more basic groups, the lower the PPB). Also, the presence of five-membered nitrogen heterocyclic rings, saturated polycyclic rings, carbonyl groups, hydroxyl groups and carboxyl groups appeared to play an important role in the reduction of PPB. (2) In high-PPB compounds (see Figure 7D-F), the presence of aromatic rings, halogen atoms (F, Cl, Br) in a benzene ring or alkyl chain, alkyl chains, sulfonyl groups, thiazoles, oxazoles and oxadiazoles etc. were associated with higher PPB values. More details are shown in Supporting Material Figure S3. and Figure S4.

**Figure 7.**
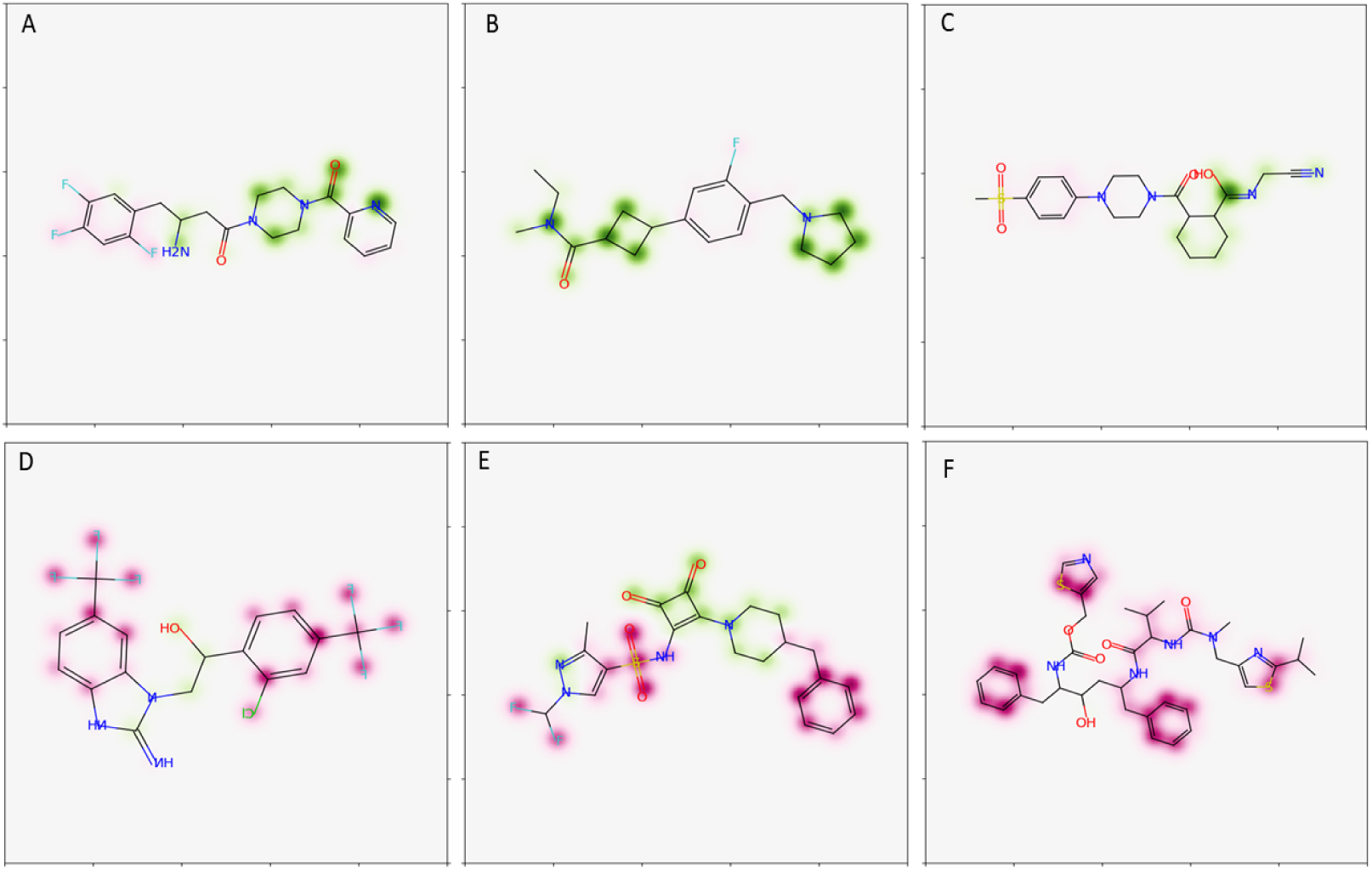
Structural contribution diagrams of partially representative low(A, B and C) and high (D, E, F) PPB compounds. Atoms colored red have a greater likelihood of increasing PPB, while green atoms are associated with lower PPB values.

We further analyzed the frequency of occurrence of substructures of high/low PPB compounds using SetCompare tool in the OCHEM platform. It was found that several functional groups were more strongly associated with one of the two analyzed classes (see Figure 8a). For example, aromatic and halogen derivatives occurred significantly more often in high PPB compounds. Additionally, further functional groups, e.g., arenes and aryl chlorides derivatives, were overrepresented in the high PPB dataset. At the same time, primary amines and secondary aliphatic amines occurred more often in the low PPB dataset (see Figure 8b). Other groups associated with low PPB are secondary alcohols, heterocycles,α, β-Unsaturated carboxylic acids and tetrahydrofurans. The results from this analysis can provide useful suggestions for chemists seeking to design compounds with lower PPB. The full list of calculated groups is show in Supporting Material.

**Figure 8.**
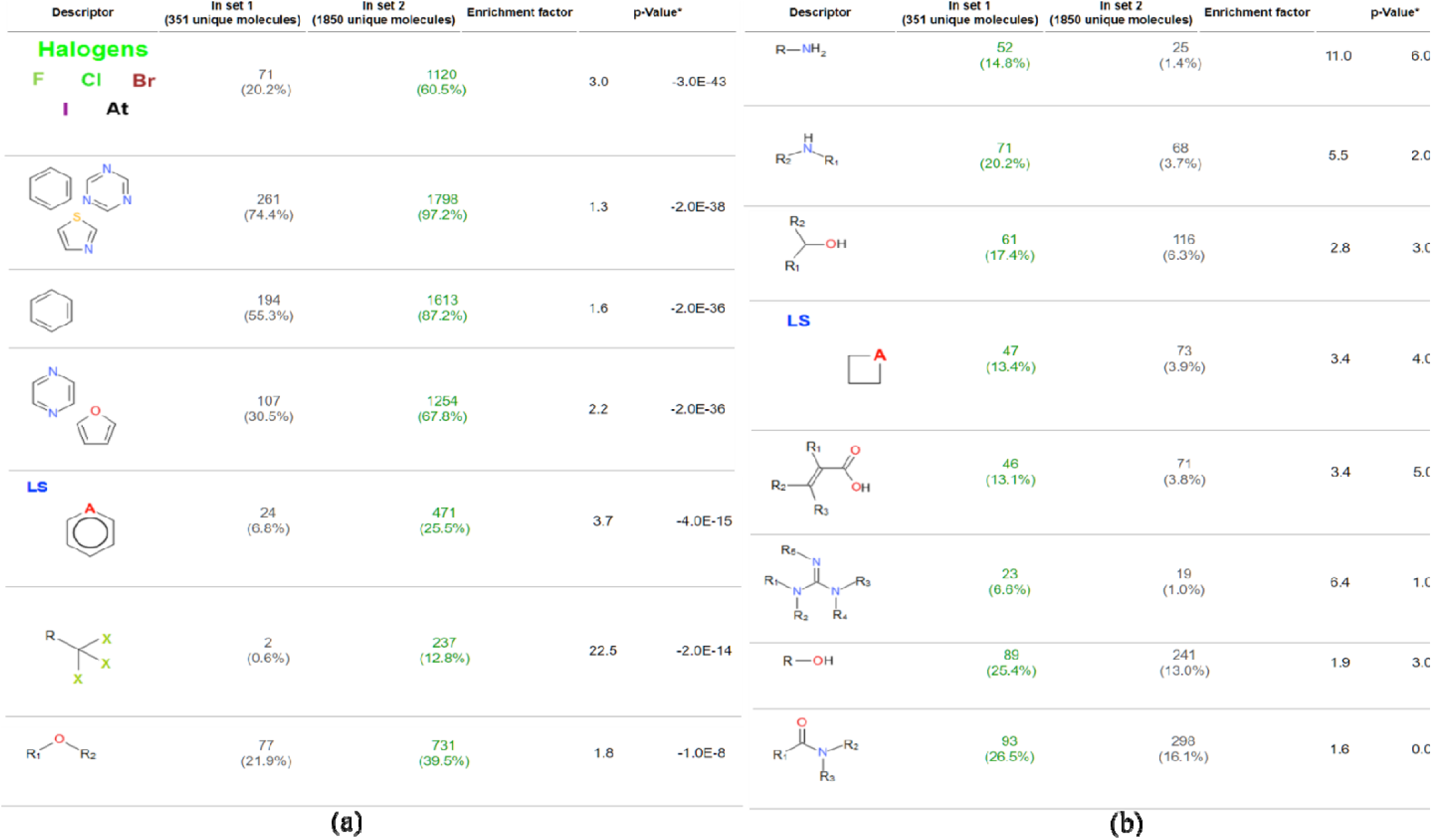
Functional groups that are overrepresented in high (a) / low (b) PPB compound datasets.

Appearance counts are listed, as well as the p-value of the respective distribution. Negative and positive p-values indicate groups overrepresented in high/low PPB datasets, respectively.

## 5. Conclusions

PPB is one of the key parameters for evaluating the efficacy of potential drugs, and the accurate prediction of PPB plays an important role in the screening stages of drug discovery. We compiled a comprehensive collection of PPB values for known compounds, rigorously curated the data, trained a variety of ML models with OCHEM and built a consensus model based on the best performing models. The consensus model performed well in both internal and external validation, and yielded results that were superior to those given by other online webservers for PPB prediction in the prospective study, i.e., predicting PPB values of diverse, unknown compounds, with experimental validation. The consensus model is available for free on the OCHEM website at https://ochem.eu/article/29. In addition, we analyzed and identified the physicochemical descriptors and fragments that have a significant influence on PPB, which may facilitate lead optimization or drug development. These results of this study are particularly useful for the prediction of high PPB compounds (>99%), as excessively high PPB poses many problems in drug discovery, such as: narrow treatment window, obvious individual differences, drug-drug interactions, and the need for clinicians to precisely measure drug concentrations. Hence, we can conclude that the current maximum achievable accuracy of PPB predictions for high PPB compounds provided by existing models is generally not satisfactory. While model described in this article demonstrated good performance for such compounds, large number of experimental measurements are required to further improve models for compounds with high PPB.

## Supporting information

The distribution of original %PPB values, ChemDiv library of 25 compounds, detection methods with HPLC or LC/MS/MS, experimental and predicted results

model output by six online web servers for 25 new measured dataset, PFC dataset, PFC99 dataset, and substructure feature analysis by Setcompare result

## Supporting Information

The distribution of original %PPB values, ChemDiv library of 25 compounds, detection methods with HPLC or LC/MS/MS, experimental and predicted results. Mordred descriptors determined via feature selection, differences in molecular and substructural properties between high and low PPB chemicals (Supporting Material.docx); model output by six online web servers for 25 new measured dataset, PFC dataset, PFC99 dataset, and substructure feature analysis by Setcompare results (Supporting Material.xlsx).

## Author Contributions

J.X. led the project, with the support from S.W. and I.V.T. Z.H. performed data collection and curation, and built ML models. Z.H., I.V.T., J.X. and Z.X jointly performed the data analysis. Z.H. tested the wet verification. Z.H. and J.X. wrote the manuscript. All authors reviewed the final version of the manuscript and provided important revisions.

## Notes

The authors declare no competing financial interest.

## Acknowledgements

We thank Dr. Katya Ahmad for proof-reading the manuscript for English and her comments.

