## Supplementary material for "The state-of-the-art machine learning model for Plasma Protein Binding Prediction: computational modeling with OCHEM and experimental validation": The distribution of original %PPB values, ChemDiv library of 25 compounds, detection methods with HPLC or LC/MS/MS, experimental and predicted results

*^a^State Key Laboratory of Bioactive Substance and Function of Natural Medicines, Institute of Materia Medica, Chinese Academy of Medical Sciences and Peking Union Medical College, Beijing 100050, China; ^b^Institute of Structural Biology, Helmholtz Munich - German Research Center for Environmental Health (GmbH), Ingolstädter Landstraße 1, 85764 Neuherberg, Germany;* ^c^*BIGCHEM GmbH, Valerystr. 49, 85716 Unterschleißheim, Germany*

**corresponding authors*

Dr. Igor V. Tetko

Institute of Structural Biology, *Helmholtz Munich - German Research Center for Environmental Health*, Ingolstädter Landstraße 1, 85764 Neuherberg, Germany

cBIGCHEM GmbH, Valerystr. 49, 85716 Unterschleißheim, Germany

Dr. Jie Xia and Song Wu

Institute of Materia Medica, Chinese Academy of Medical Sciences,

No. 2 Nanwei Road, Beijing 100050, China

**Figure S1. The distribution of original %PPB values**

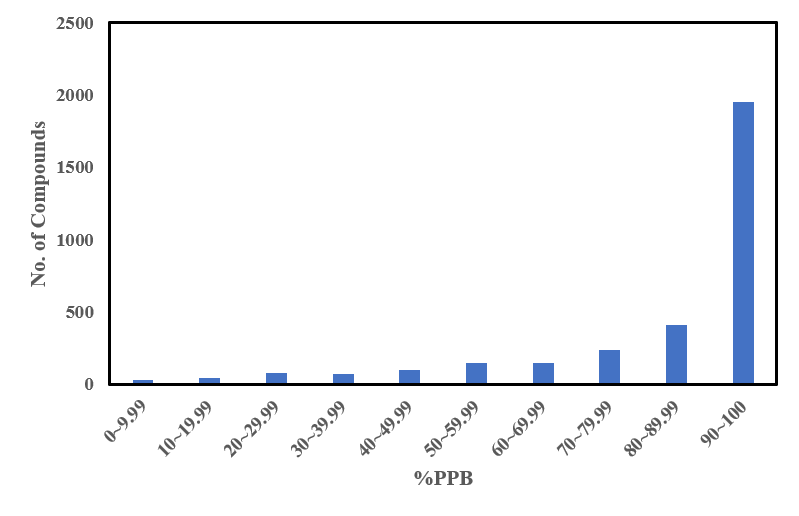

**Table S1. ChemDiv library of 25 compounds HPLC or LC-MS/MS detection methods**

| ID | SMILES | Mobile phase A% | Mobile phase B% | UV(nm) |
| --- | --- | --- | --- | --- |
| S941-0208 | O=C(N1CC(CC2=NN=CN2C3CCCC3)C1)C=4C=CSC4 | 80 | 20 | 245 |
| L498-0418 | COC1=CC=CC(NC(=O)C2=CC=C(C=C2)C=3ON=C(N3)C4=CC=C(F)C=C4)=C1 | 40 | 60 | 246 |
| E589-3042 | CC1CC2=CC(=CC=C2N1C(=O)C)S(=O)(=O)NCC3CCC(CC3)C(=O)N4CCCCC4 | 60 | 40 | 271 |
| C390-0229 | CC1=CC=C(C=C1)N2C(C3=CC=CC(=C3)[N+](=O)[O-])C=4C(C)=NNC4C2=O | 60 | 40 | 252 |
| D674-0084 | CC1=CC=CC=C1NC(=O)COC2=CC=CC(=C2)C3=NOC(=N3)C4=CC=CC=C4Cl | 40 | 60 | 243 |
| L707-1990 | COC1=CC=CC(NC(=O)CN2C(=O)CCCC3=CC(=CC=C23)S(=O)(=O)N4CCCCCC4)=C1 | 50 | 50 | 248 |
| F855-0012 | CCC(C)N1N=NC2=CC(=CC=C12)S(=O)(=O)N3CCC(CC3)C(=O)NCC4=CC=CC=C4Cl | 50 | 50 | 270 |
| G799-0300 | CCOC(=O)C=1NC(C)=C(CN(C2CCCCC2)C(=O)NC3=CC=CC(SC)=C3)C1C | 40 | 60 | 241 |
| D134-0413 | COC1=CC=C(C=C1)C2=NOC(CCCC(=O)NCC3=CC=CC=C3)=N2 | 50 | 50 | 258 |
| E958-1454 | COC1=CC=C(CCNCC2=CC=CN2C3=NN=C(S3)N4CCCCCC4)C=C1OC | 75 | 25 | 308 |
| 2279-4329 | CC(NC1=NN=C(O)N=C1O)C(=O)NN=CC2=CC=CC(C)=C2 | 75 | 25 | 283 |
| D400-0405 | CCC1=NOC(=N1)C2=CC=CC(OCC(=O)NC3=CC=CC(C)=C3)=C2 | 40 | 60 | 246 |
| L711-0358 | CC1=CC=C(C=C1)N2N=C(C)C=3SC(=NC23)N4CCCC(C4)C(=O)NC5CCCCC5 | 40 | 60 | 278 |
| J075-4126 | BrC1=CC=C(C=C1)C(=O)NCC2=NOC(=N2)C3=CC=CC=C3 | 50 | 50 | 246 |
| K284-3537 | FC1=CC=CC=C1CSC2=NC3=CC=CC=C3C(=O)N2CCCC(=O)NCC=4OC=CC4 | 40 | 60 | 276 |
| 8012-8788 | FC1=CC=CC(=C1)C=2OC(=NN2)C3=CC=C(OCC4=CC=CC(Br)=C4)C=C3 | 30 | 70 | 298 |
| D360-5516 | CC1=CC=C(NC(=O)CSC2=NC=3CCCCC3C=C2C#N)C=C1C | 40 | 60 | 266 |
| L009-0033 | COC1=CC=CC(NC(=O)C2=CC=CC(=C2)C3=CC=C(N=N3)N4CCCC4)=C1 | 80 | 20 | 270 |
| K072-0227 | CN(N=CC1=CC=CN=C1)C2=C(Cl)C(=O)N(N=C2)C3=CC=C(F)C(Cl)=C3 | 70 | 30 | 358 |
| D294-5147 | COC1=CC=CC=C1CNC(=O)C2CN(CCC3=CNC4=CC=C(F)C=C34)C(=O)C2 | 60 | 40 | 220 |
| C878-1846 | CC1=CC=CC(CSC2=NC3=CC=NC=C3N2CC4=CC=C(C=C4)C(=O)NCC5=CC=C(F)C=C5)=C1 | 70 | 30 | 240 |
| 2034-0020 | CC1=NN(C(=O)C1=CC2=CC=C(Cl)C=C2Cl)C3=CC=CC=C3 | 40 | 60 | 245 |
| S732-0033 | O=C(N1CCCCCCNC(=O)C2CCCC12)C3=CC=CC(=C3)C#N | 60 | 40 | 220 |
| 4984-5643 | CCOC1=CC(=CC=C1OCC2=CC=C(F)C=C2)C3C4=C(CCCC4=O)NC5=C3C(=O)CCC5 | 50 | 50 | 242 |
| D087-0017 | O=S(=O)(C1CCS(=O)(=O)C1)N2CCCC2 | LC-MS/MS analysis | | |

The HPLC system was composed of a Shimadzu LC-20AT and DAD detector (Shimadzu Scientific Instruments, Tokyo, Japan) liquid chromatograph. The analytical column was an Inertsil ODS-3 (150*3.0mm, 3μm). The mobile phase consisted of water containing 0.1% phosphoric acid (phase A) and acetonitrile (phase B) at a flow rate of 0.7 ml/min with an operating temperature at 40℃. The elution procedure is isocratic and the detection time of each sample does not exceed 20 min. The chromatographic conditions for each sample are shown in Table 1.

Compound 25 has no UV absorption, so it was detected using LC-MS/MS methods. The LC–MS/MS system was composed of a Agilent 1290 Infnity and 6495C mass spectrometry (Agilent Technologies, USA). Detection was carried out on a triple quadrupole tandem mass spectrometer equipped with electrospray ionization (ESI) in positive ion multiple reaction monitoring (MRM) mode. The compound 25 were separated on an a Zorbax Eclipse Plus C18 (50*2.1mm, 1.8μm) column by two mobile phases: mobile phase A (0.1% formic acid in water [v/v]) and mobile phase B (0.1% formic acid in acetonitrile [v/v]), and was operated with a gradient elution at 0.4 mL/min as follows: 90% A (0–0.5 min), 90% A → 25% A (0.5–3.0 min), 25% A → 90% A (3.0–3.01 min), 90% A (3.01–4.0 min). The total run of the analysis took 4 min. The optimized Nozzle Voltage and capillary were set at 500V and 4000V. Nitrogen was used as sheath gas and flow and temperature at 11L/min and 250℃, nebulizer pressure at 20psi, nitrogen was used as drying gas and flow and temperature at 14L/min and 200℃. The collision gas (high purity nitrogen) pressure was 0.15Mpa and the collision energy were 13 Ev for compound 25. The MRM transitions monitored were 254.0→190.0, 254.0→136.0 for quantitative and qualitative, respectively.

**Table S2.** **Mordred descriptors determined via feature selection and Pearson correlation coefficient (R^2^) between the descriptor and PPB.**

| **Descriptors** | **R^2^** | **Description** |
| --- | --- | --- |
| SLogP | 0.604739 | Wildman-Crippen LogP |
| FilterItLogS | 0.588603 | Filter-it™ LogS |
| C2SP2 | 0.499323 | SP2 carbon bound to 2 other carbons |
| ETA_beta_ns | 0.490627 | nonsigma contribution to valence electron mobile count |
| SMR_VSA7 | 0.485139 | MOE MR VSA Descriptor 7 ( 3.05 <= x < 3.63) |
| nAromAtom | 0.484390 | nAromAtom |
| piPC3 | 0.474827 | 3-ordered pi-path count (log scale) |
| NaasC | 0.467597 | number of aasC |
| piPC10 | 0.457000 | 10-ordered pi-path count (log scale) |
| TpiPC10 | 0.455782 | 10-ordered total pi-path count (log scale) |
| ETA_dEpsilon_D | 0.450276 | ETA delta epsilon (type: D) |
| SlogP_VSA6 | 0.446229 | MOE logP VSA Descriptor 6 ( 0.15 <= x < 0.20) |
| ETA_beta | 0.440862 | valence electron mobile count |
| piPC1 | 0.436051 | 1-ordered pi-path count (log scale) |
| NaaCH | 0.419311 | number of aaCH |
| AATS1v | 0.411642 | autocorrelation of lag 1 weighted by vdw volume |
| ZMIC1 | 0.402523 | 1-ordered Z-modified information content |

**Figure S2. The** **differences in molecular properties between high and low PPB chemicals**

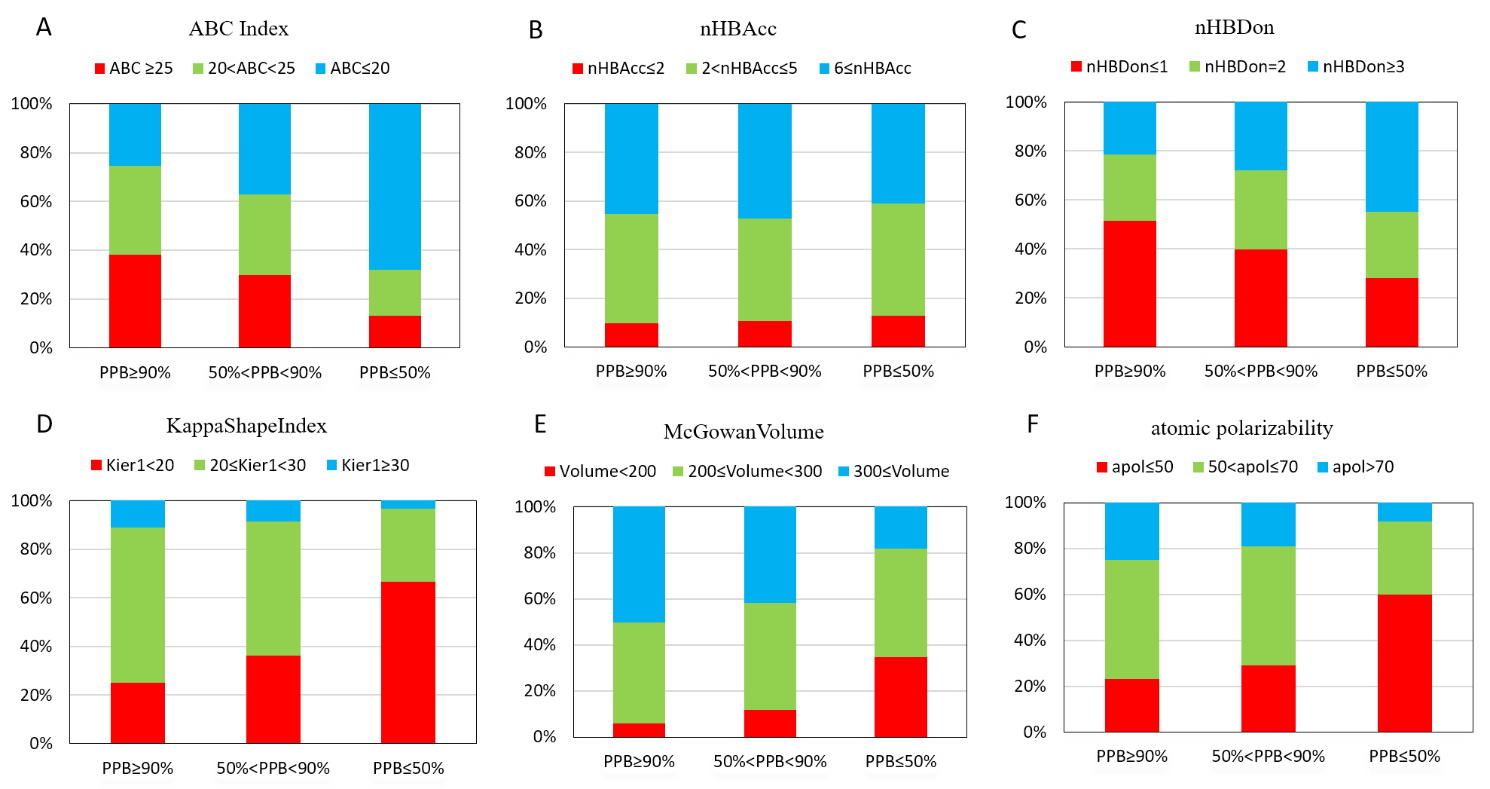

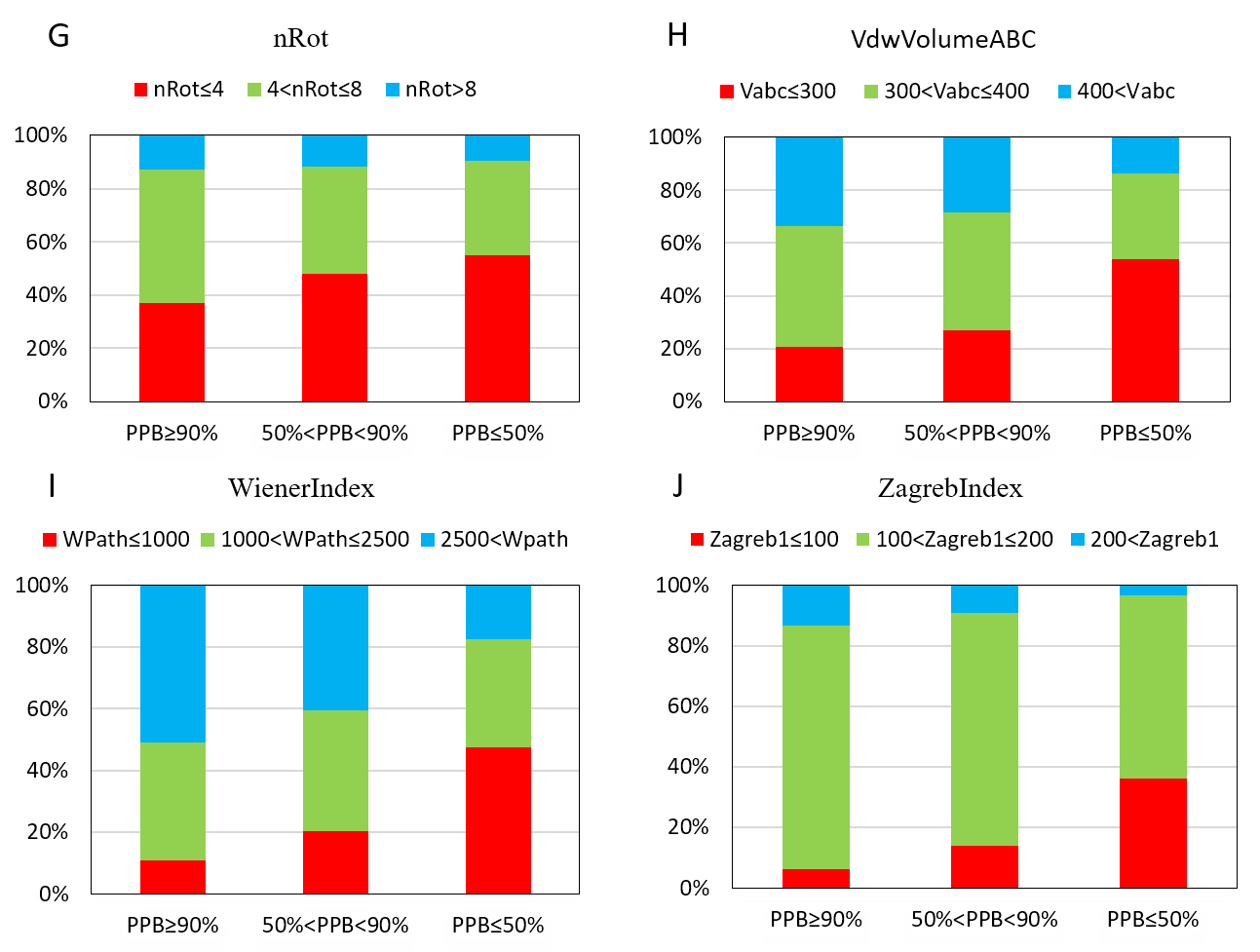

**Figure S3. Low-PPB compounds** **substructural feature**

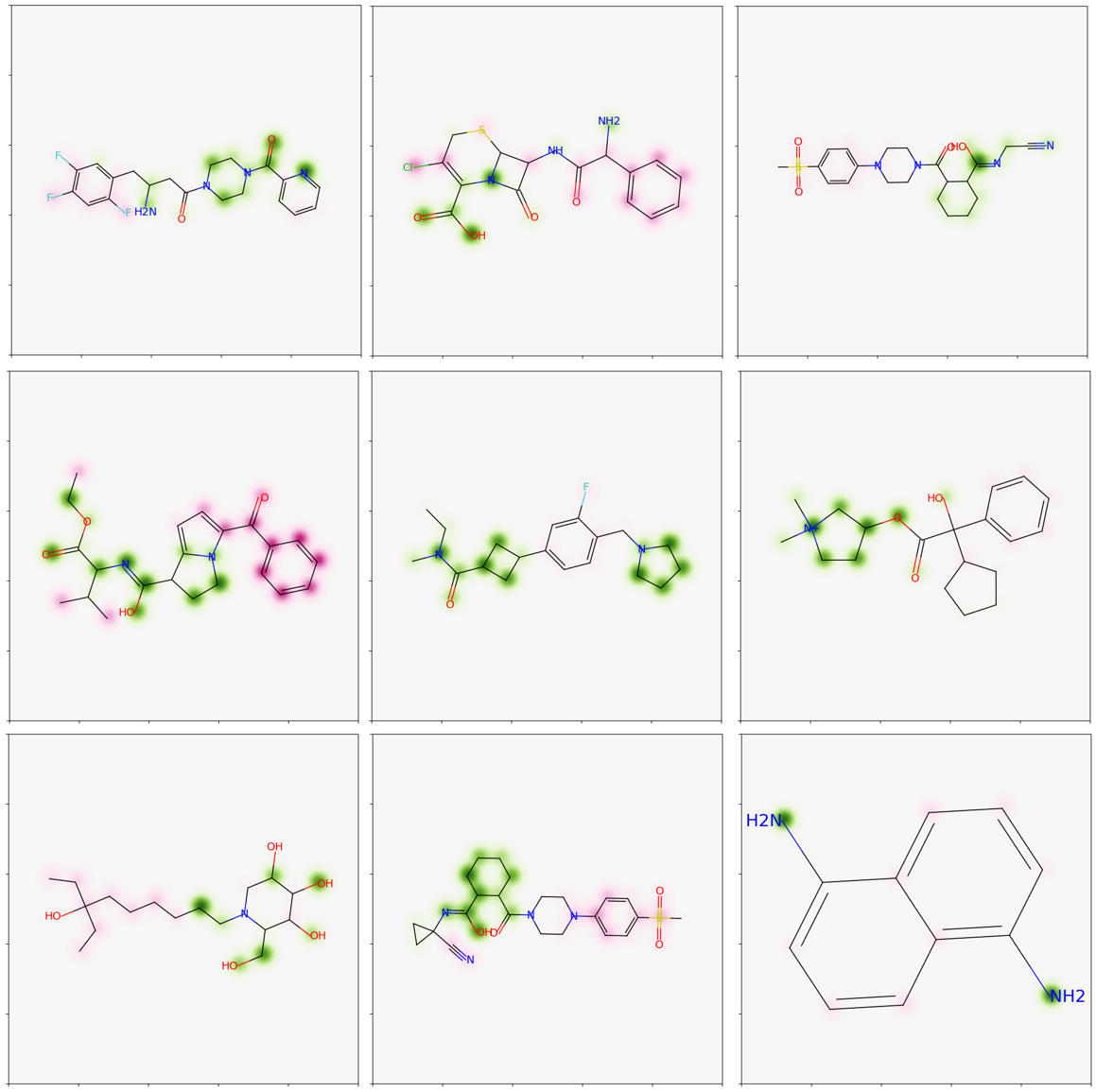

**Figure S4. High-PPB compounds substructural feature**

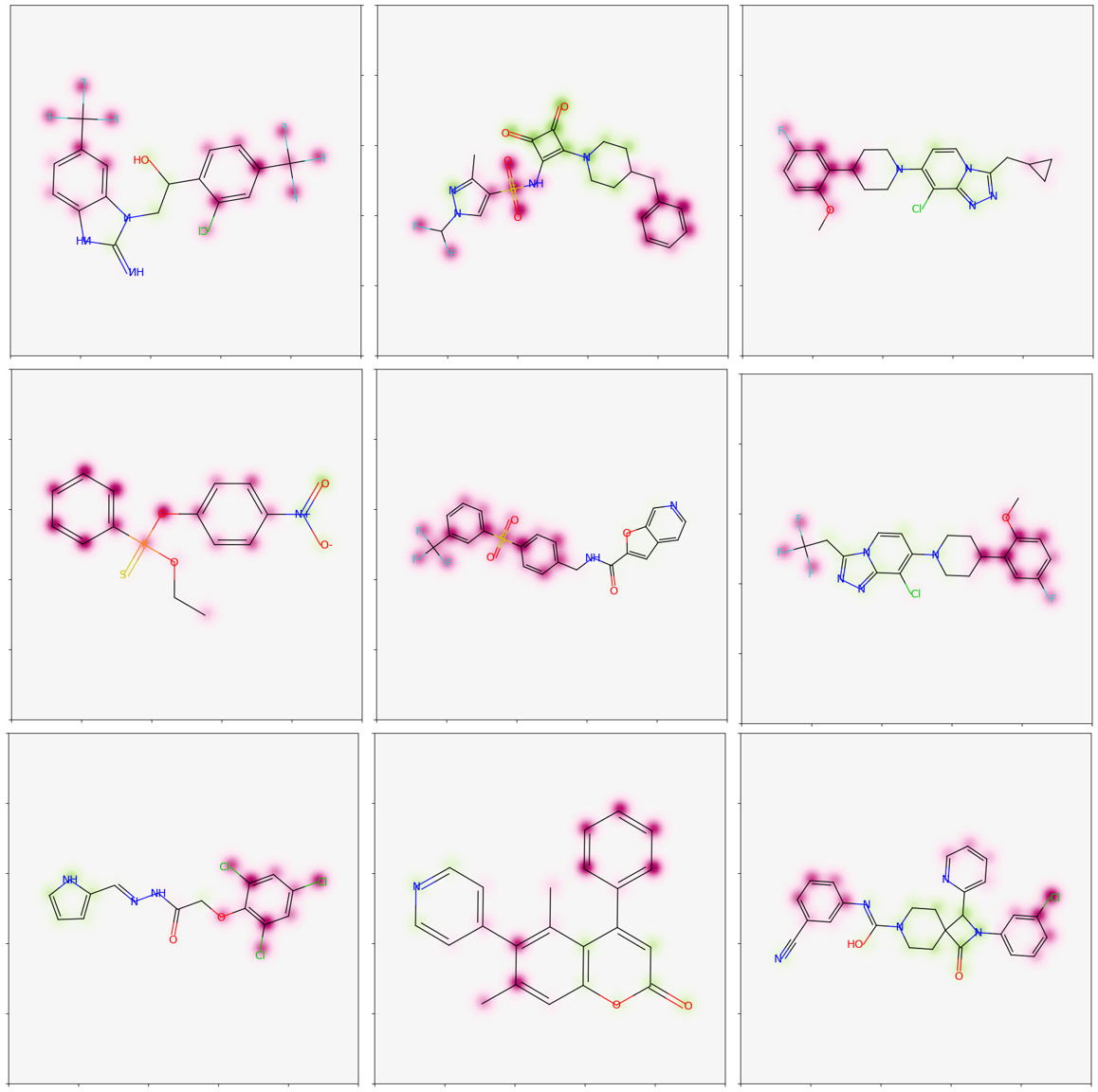
